# Meiotic recombination dynamics in plants with repeat-based holocentromeres shed light on the primary drivers of crossover patterning

**DOI:** 10.1101/2023.04.28.538594

**Authors:** Marco Castellani, Meng Zhang, Gokilavani Thangavel, Yennifer Mata Sucre, Thomas Lux, José A. Campoy, Magdalena Marek, Bruno Huettel, Hequan Sun, Klaus F. X. Mayer, Korbinian Schneeberger, André Marques

## Abstract

Centromeres strongly affect (epi)genomic architecture and meiotic recombination dynamics influencing the overall distribution and frequency of crossovers. Here, we studied how recombination is regulated and distributed in the holocentric plant *Rhynchospora breviuscula*, a species lacking localised centromeres. Combining immunocytochemistry, chromatin analysis and high-throughput single-pollen sequencing, we discovered that crossover frequency is higher at ends related to centred chromosomal regions. Contrasting the diffused distribution of (epi)genetic features and hundreds of repeat-based centromeric units. Remarkably, we found that crossovers were abolished at core centromeric units but not at their vicinity indicating the absence of a centromere effect across repeat-based holocentromeres. We further show that telomere-led pairing and synapsis of homologous chromosomes appear to be the primary force determining the observed U-shaped recombination landscape. While centromere and (epi)genetic properties only affect crossover positioning locally. Our results suggest that the conserved U-shaped crossover distribution of eukaryotes is independent of chromosome compartmentalisation and centromere organisation.

## Introduction

During meiosis, homologous chromosomes undergo meiotic recombination, in which genomic material is exchanged between homologous chromosomes. This exchange is initiated by the physiologically induced DNA double strand breaks (DSBs)^1, 2^. The formation of meiotic DSBs is commonly resolved via crossovers (COs) as well as other recombination outcomes, referred to as non-COs (NCOs)^3^.

Crossovers can be divided into two classes, although the existence of alternative CO pathways cannot be excluded^4, 5^. Class I COs are the most prevalent and are sensitive to interference, i.e., they do not occur near each other along a chromosome. Class I COs result from the ZMM pathway that includes the key factor Human enhancer of invasion-10 (HEI10), involved in CO designation, and ZYP1, a key protein involved in synaptonemal complex (SC) assembly^6–12^. Class II COs are insensitive to interference and accommodate around of 10% of the total COs in *Arabidopsis thaliana*^13^.

The global distribution of COs is typically associated with the distribution of genetic and epigenetic [(epi)genetic] features^14, 15^. In most eukaryotes, gene/euchromatin density positively correlates with CO frequency^16, 17^. By contrast, CO frequency is typically lower in heterochromatic regions, including at (peri)centromeres^18, 19^. In monocentric species, centromeres are single, defined structural entities and are typically repeat-based. Recombination is largely suppressed at and in the proximity of centromeres in these species, a phenomenon known as the centromere effect^20^. In plants with large chromosomes the centromere effect can extend several megabases (Mb) along pericentromeric regions, that can represent a large proportion of the chromosomes^21–23^. Monocentricity is not the only centromeric organisation adopted by eukaryotes, however. For instance, holocentric species harbour multiple centromeric determinants over the entire length of their chromosomes^24, 25^. Thus, it would be interesting to understand how COs are regulated in holocentric species where hundreds of centromeric units are distributed chromosome-wide.

Holocentricity has evolved independently multiple times during the evolution of nematodes, insects and plants^26, 27^. In the holocentric animal models *Caenorhabditis elegans* and silk moth (*Bombyx mori*), holocentromeres do not associate with a specific sequence and thus can have cell-specific dynamics^28, 29^. By contrast, holocentric plants of the *Rhynchospora* genus (beaksedges) display repeat-based holocentromeres in both mitosis and meiosis^30, 31^. Recently, we sequenced the genomes of three beaksedges (*R. breviuscula*, *R. pubera* and *R. tenuis*) and determined that each chromosome harbours multiple short arrays (∼20 kb each) of the specific *Tyba* tandem repeat, evenly spaced (every 400–500 kb) along the entire chromosomal length, and specifically associated with centromeric histone H3 protein CENH3^32^. This particular chromosome organisation is associated with remarkably uniform distribution of genes, repeats, and epigenetic features, in stark contrast to the compartmentalised chromosome organisation of close monocentric relatives^32^. Remarkably, each individual centromeric unit in *R. pubera* showed very similar epigenetic regulation as found in other plant monocentromeres^32, 33^. Thus, beaksedges offer an excellent model to study the mechanisms of CO formation in the absence of the major effect of the monocentromere, while having similar centromere chromatin (epi)genetic properties.

Regardless of being monocentric or holocentric, most studied eukaryotes show higher recombination rates at distal chromosomal arm regions. This typical U-shape CO distribution is usually explained as the result of structural chromosome features (telomere and centromere effects) and correlation with (epi)genetic factors^14, 17, 33–36^. However, how these factors specifically influence meiotic recombination patterning at broad and local scales are still not known. Understanding the uniform distribution of (epi)genetic features and absence of conventional centromeres in *Rhynchospora* will allow us to explore conserved and adapted mechanisms influencing meiotic recombination patterning among eukaryotes. Studies of meiosis in holocentric plants have been mainly focused on the intriguing phenomenon of “inverted meiosis”^26, 37–39^. Moreover, meiotic chromosomes in *Rhynchospora* maintain the repeat-based holocentromere organisation^31^, suggesting that COs can be formed very close to centromere chromatin. No direct evidence of meiotic recombination frequency and distribution has yet been reported for any holocentric plant. It is still unknown whether and how plant holocentromeres interact or interfere with meiotic recombination.

Here, we use *R. breviuscula* as a model to study meiotic recombination dynamics in the absence of both a localised centromere and a compartmentalised chromosome organisation, features that potentially mask underlying factors affecting the CO distribution in most eukaryotic genomes. Using a combination of immunocytochemistry, chromatin and DNA analysis, and CO calling from high-throughput single-pollen sequencing, we develop a comprehensive overview of meiotic recombination dynamics and distribution for a species with repeat-based holocentromeres. We show that despite this unique chromosome organisation, COs distribution is biased towards the distal regions of chromosomes, forming a typical U-shape distribution. Importantly, this distribution did not correlate with any (epi)genetic feature analysed at broad scale. Remarkably, we found that COs are suppressed at core repeat-based centromeric units but not at their vicinity, indicating the absence of a centromere effect. We show that chromosome ends have higher CO frequency even in the absence of a monocentromere and compartmentalised (epi)genomic features. In fact, our data suggest that pairing and synapsis dynamics starting from chromosomal ends exert a major influence in determining the broad-scale recombination landscape, whether a centromere is present or not. We propose that centromere and (epi)genetic features play a role in CO positioning only at fine-scale.

## Results

### The molecular dynamics of meiosis I is conserved in *R. breviuscula*

Chromosome spreads on male meiocytes of *R. breviuscula*, allowed us to conclude that prophase I progression is conserved in this species. We observed all the classical prophase I stages, e.g., leptotene, zygotene, pachytene, diplotene, and diakinesis (**Figure S1A–E**). In contrast to the holocentric animal *C. elegans* ^35^, which forms only a single chiasma per bivalent, we observed the presence of five bivalents connected by one or two chiasmata in *R. breviuscula* (**Figure S1E**), consistent with reports in other holocentric plants^39^. Moreover, we confirmed the holocentric nature of *R. breviuscula* chromosomes in mitosis and meiosis by showing the localisation of the centromeric protein CENH3 (**Figure S1E–H**).

We then investigated the immunolocalisation of ASY1^40, 41^ and ZYP1^6, 7, 11, 12, 42^ as indicators of a conserved and functional machinery for chromosome axis and synapsed regions, respectively. The ASY1 signal was present along the entire length of unsynapsed chromosomes in early prophase I, corresponding to leptotene (**Figure 1A**). During zygotene, the SC started to assemble and ZYP1 was gradually loaded onto synapsed chromosomes. As ZYP1 was loaded, the two ASY1 linear signals could be followed until they converged and lost intensity, after which the ZYP1 linear signal became clear and intense (**Figure 1B,C**). As meiosis progressed into pachytene, with complete synapsis and pairing, we detected the linear ZYP1 signal along the full length of chromosomes (**Figure 1D**). The ZYP1 signal localised in the groove between the pairs of homologous chromosomes. The combined behaviour of ASY1 and ZYP1 was consistent with that observed in monocentric models. This hints at a conserved pairing and synapsis mechanism in *R. breviuscula*, despite the CENH3 distribution along the entire length of synapsed chromosomes during meiosis (**Figure S1H**). We also tested whether the meiosis-specific alpha-kleisin REC8 is also conserved in *R. breviuscula*. REC8 is responsible for sister chromatid cohesion and is important for chromosome segregation and recombination^43^. Indeed, we detected a conserved linear REC8 signal at pachytene, when REC8 co-localised with ZYP1 as a continuous linear signal along the entire synapsed chromosomes (**Figure 1D**). Thus, pairing and synapsis are conserved in the holocentric plant *R. breviuscula*, resembling those in monocentric models.

**Figure 1.**
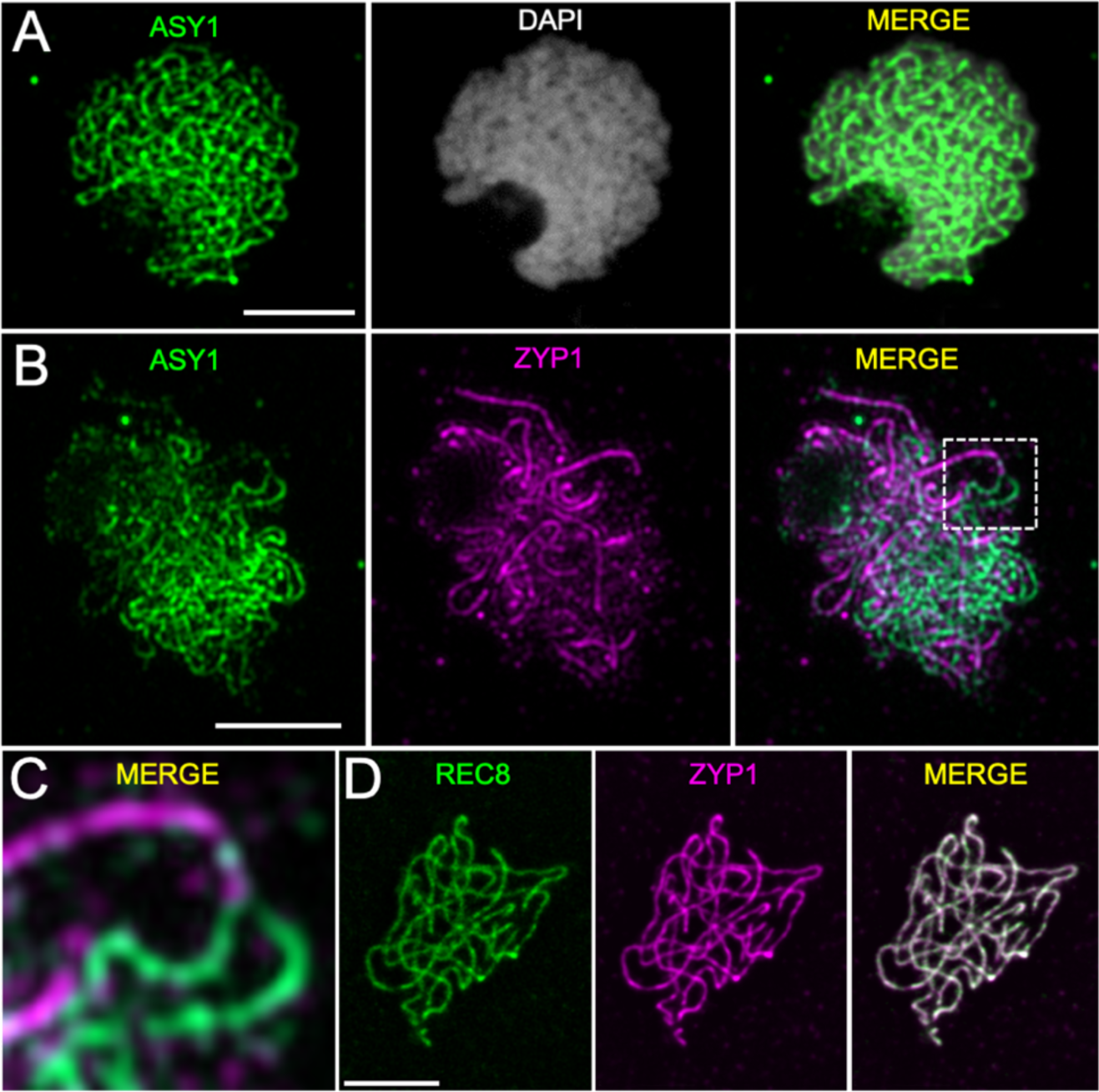
Immunolocalisation of ASY1, REC8, and ZYP1 from leptotene to pachytene of meiosis prophase I. **(A)** ASY1 (green) appears as a linear signal on unpaired chromosomes. **(B)** Synapsis is visualised as the loading of ZYP1 (magenta) as the ASY1 (green) signal disappears. **(C)** Magnification of two unpaired chromosomes (dashed square in **B**), represented by ASY1 (green), coming together to synapse, with the loss of the ASY1 signal and loading of ZYP1 (magenta). **(D)** Full co-localisation of cohesin protein REC8 (green) and ZYP1 (magenta) at pachytene. A maximum projection is shown; chromosomes were stained with DAPI. Images were acquired with a Zeiss Axio Imager Z2 with Apotome system and Leica Microsystems Thunder Imager Dmi8 (Scale bar of detail in **C**, 2 µm; other scale bars, 5 µm).

We further studied the behaviour of HEI10, a RING-family E3 ligase that has been characterised in mammals, yeast (*Saccharomyces cerevisiae*) and plants. HEI10 functions after synapsis has occurred in ZMM pathway but before the resolution of COs. HEI10 has been proposed to interact with both early and late recombination proteins, acting by stabilising recombination sites and promoting their maturation into class I COs^8, 9, 44^. In *R. breviuscula*, when pairing and synapsis started in early zygotene, HEI10 was immediately loaded as closely spaced signals co-localising with the first ZYP1 signals (**Figure 2A**). In pachytenes HEI10 progressed to form a linear signal co-localising with ZYP1 along the entire synapsed chromosomes (**Figure 2B**). During pachytene, when synapsis is complete, the HEI10 linear signal started to become non-homogeneous along chromosomes, while a few foci increased in intensity (**Figure 2C**). We think that these are putatively class I CO sites. In diplotene and diakinesis, only the high-intensity foci remained (**Figure 2D**). Thus, the dynamic behaviour of HEI10 is conserved and most likely the recently proposed HEI10 “coarsening” model is acting similarly in *R. breviuscula*^45–47^.

**Figure 2.**
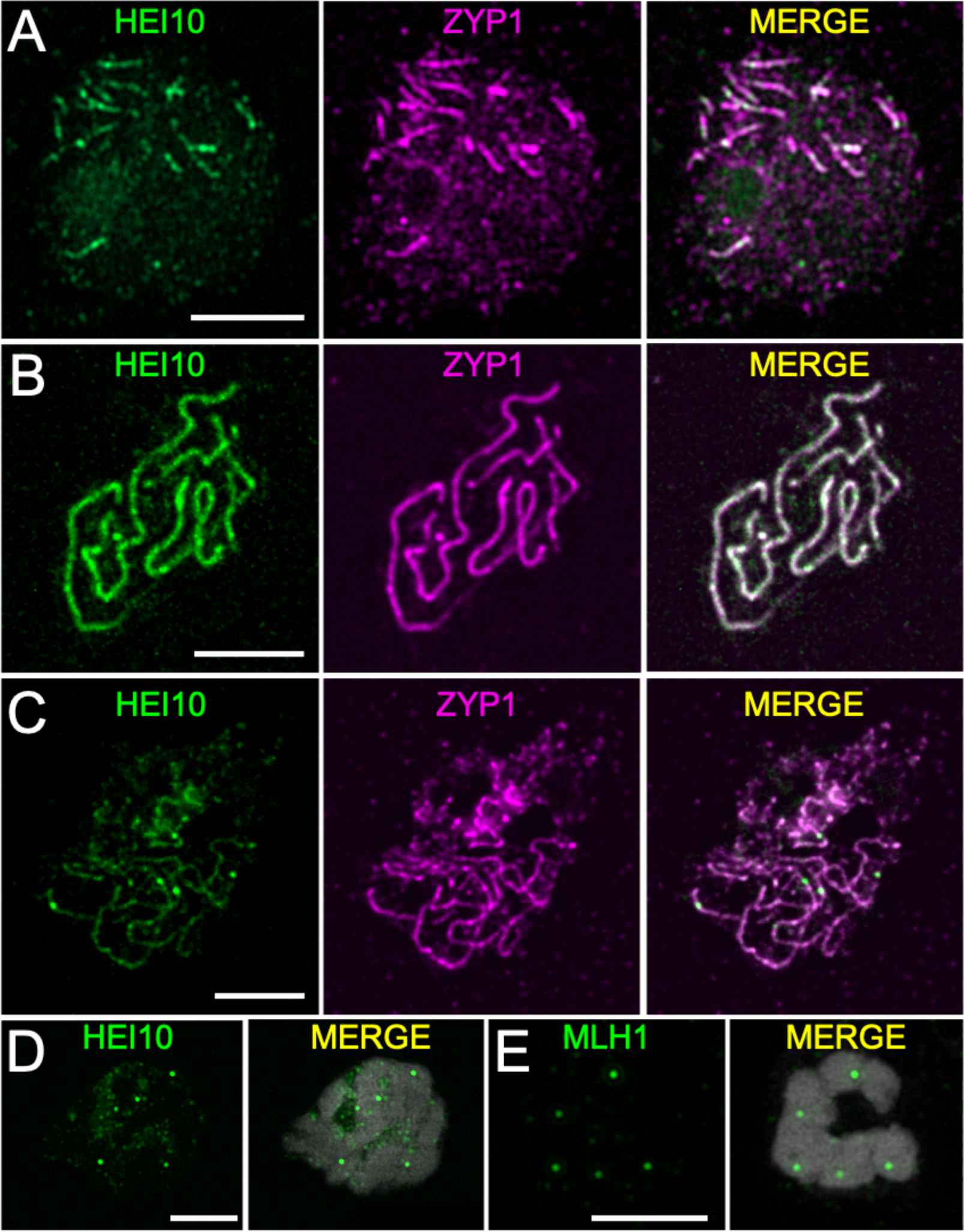
Immunolocalisation of HEI10, ZYP1 and MLH1 in late prophase I. **(A)** In early zygotene when synapsis starts, HEI10 (green) is immediately loaded as many closely spaced foci co-localising with ZYP1 (magenta). **(B)** In pachytene, HEI10 (green) is visible appearing as lines, which co-localise with ZYP1 (magenta). **(C)** In late pachytene, the linear signal of HEI10 still co-localises with ZYP1, but becomes weaker, except for a few highly intense foci. **(D)** During the diplotene and diakinesis stages, HEI10 appears only as foci on bivalents, with no linear signal. **(E)** MLH1 (green) appears in late prophase I stages as foci on bivalents, representing chiasmata. Maximum projections are shown, with chromosomes stained with DAPI. Images were acquired with a Zeiss Axio Imager Z2 with Apotome system and Leica Microsystems Thunder Imager Dmi8. Scale bars, 5 µm.

Another established marker for meiotic recombination is the mismatch repair protein MLH1 (MUTL-HOMOLOG 1), which is essential for meiosis and is believed to have a meiosis-specific resolvase activity to process double Holliday junctions (dHJs) into final class I COs. MLH1 interacts with MSH4 (MUTS HOMOLOG 4) and MSH5 in the dHJ resolution pathway, thus specifically marking class I COs in distantly related species^48^. In *R. breviuscula*, MLH1 appeared as bright foci on bivalents during the diplotene and diakinesis stages (**Figure 2E**). We detected at least five foci, one on each bivalent, with a maximum of eight foci (**Figure S2**), which is consistent with the formation of two COs in some bivalents. The mean number of foci was 6.27 (*n* = 83).

### Phased genome assembly of *R. breviuscula* as a prerequisite for CO identification by gamete-sequencing

Determining whether recombination in *R. breviuscula* is affected by the genome-wide distribution of holocentromeres requires the detection of CO events in a large number of recombinant individuals. However, *R. breviuscula* is an outbred wild species with high levels of self-incompatibility, which hampers the standard detection of COs, typically involving the time-consuming generation of segregating offspring. As gametes already carry the outcome of meiotic recombination and can be obtained in large numbers in a relatively inexpensive manner from pollen grains, we adapted a strategy based on the gamete-binning method described by Campoy et al.^49^ (see below). To identify COs from a single *R. breviuscula* individual, the genome of the given organism must be heterozygous and a phased chromosome-level reference genome must be available. The recently reported nonphased genome of *R. breviuscula* was reported to be 1% heterozygous^32^, suggesting the feasibility of haplotype phasing the genome. We took advantage of the recent development of the assembler software Hifiasm^50^, which enables the accurate phasing of both haplotypes from primary assembled contigs using a combination of HiFi reads and Hi-C (see Online Methods; **Figure 3A,B**). Further Hi-C scaffolding of each set of haplotype-phased contigs led to high-quality haplotype-phased chromosome-level genome assemblies (**Figure 3C; Table S1**). We performed a synteny analysis and detected the structural variants between the two haplotypes, revealing a high degree of synteny between the haplotypes with only few inversions, translocations and duplications (**Figure 3D; Table S2**).

**Figure 3.**
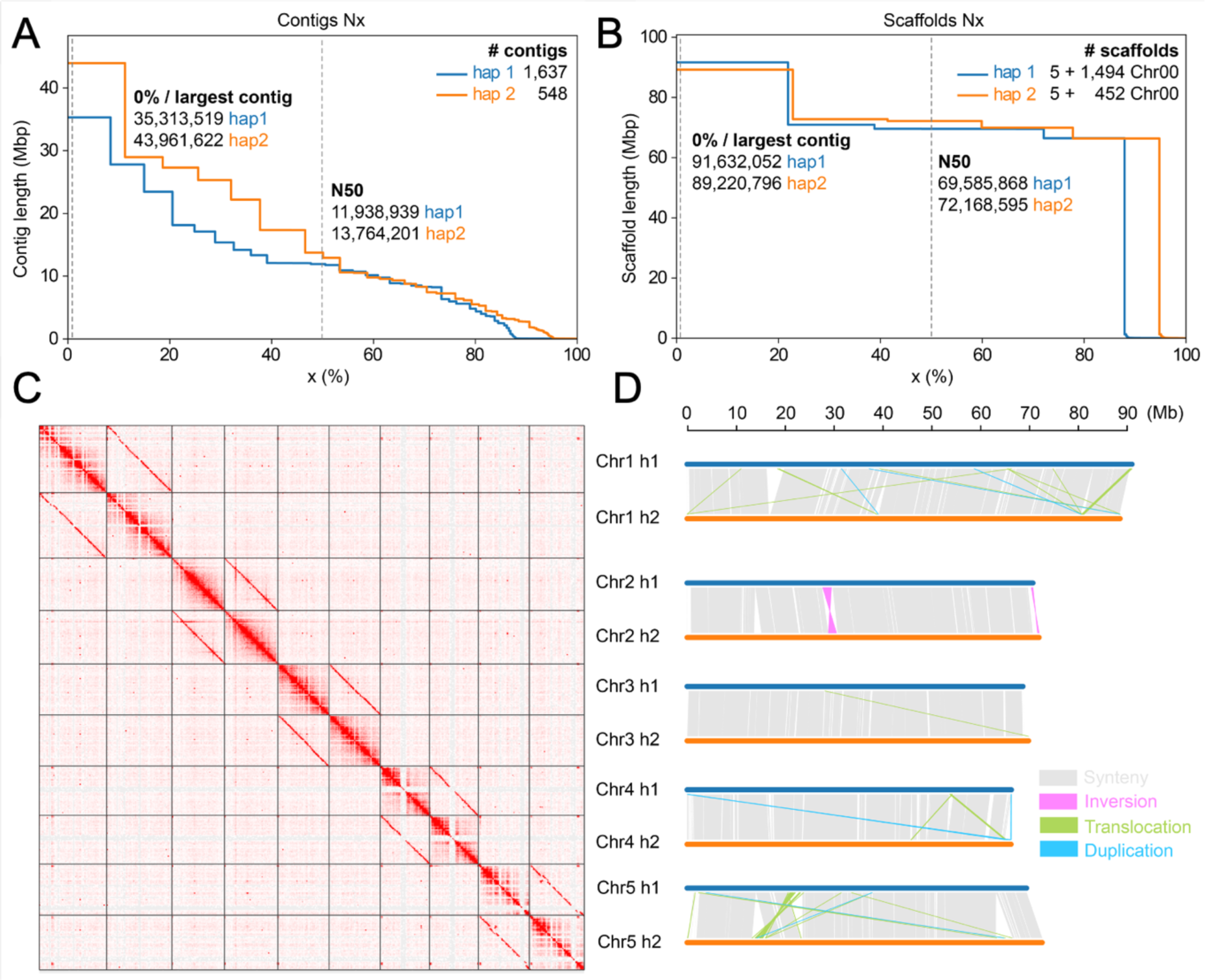
Phasing and structural variation of the *R. breviuscula* heterozygous genome. **(A,B)** Assembly statistics of the phased contigs **(A)** and scaffolds **(B)** for haplotype 1 and haplotype 2. **(C)** Hi-C scaffolding of the five haplotype-phased pseudochromosomes. Homozygous regions between the haplotypes are seen as clear regions depleted of signals on the Hi-C map. **(D)** Synteny assessment and structural variants (>10 kb) identified between the two haploid assemblies. Note the overall high synteny between the two haplotypes. Synteny blocks were computed with SyRI^51^.

To genotype the haploid gamete genomes and determine from which haplotype a genomic segment is derived, genome-wide markers are needed to distinguish the two haplotypes. By aligning the ∼26-Gb Illumina whole-genome short reads of *R. breviuscula* with the haplotype 1 phased genome (from the reference genome, rhyBreHap1), we detected 820,601 haplotype-specific single nucleotide polymorphisms (SNPs, ∼1 SNP/449 bp) and used them as markers for genotyping (**Figure 4B; Figure S3, S5A**).

**Figure 4.**
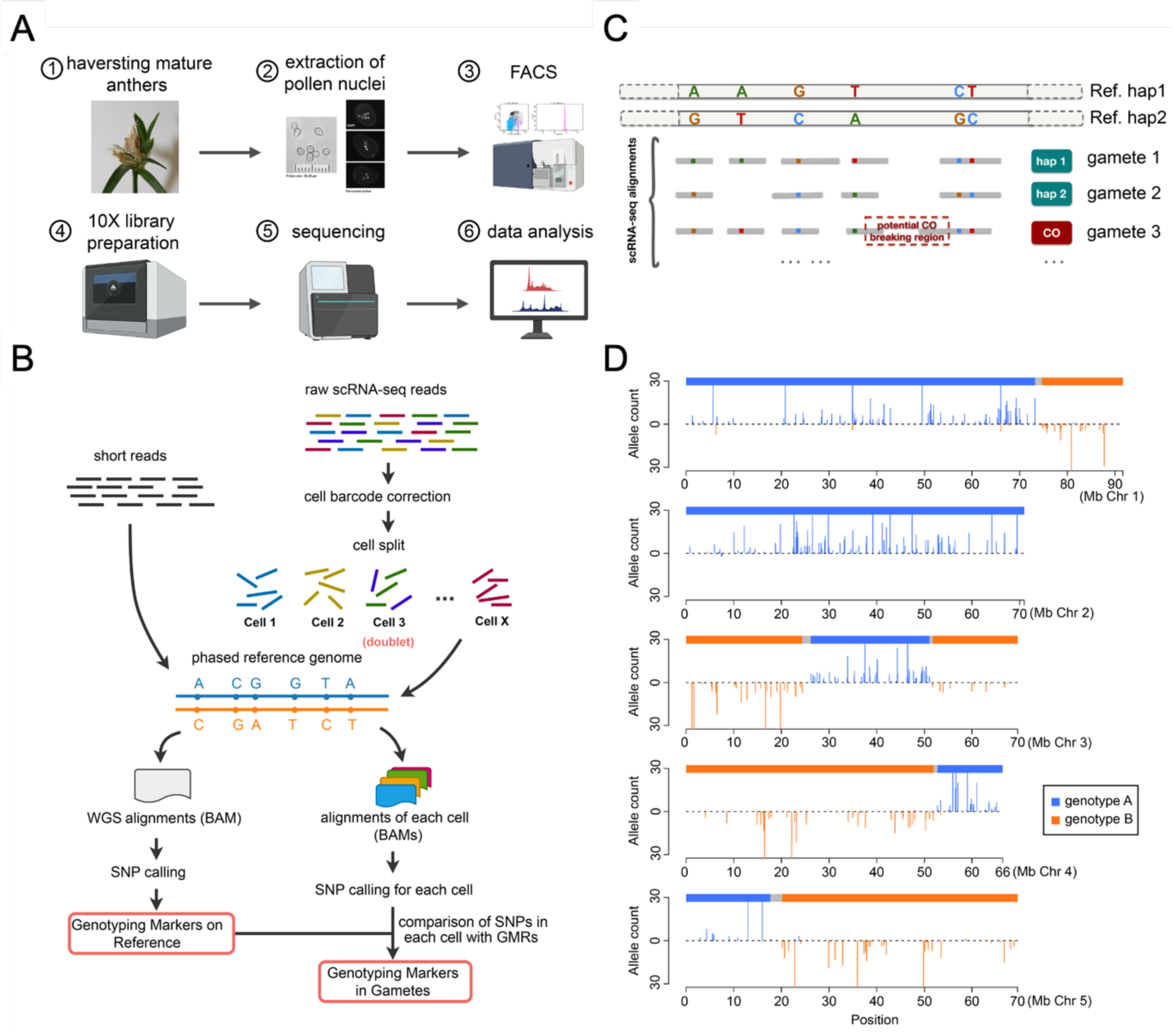
Overview of CO calling by adapting scRNA sequencing to *R*. *breviuscula* gametes. **(A)** Pollen sampling, library preparation and scRNA sequencing pipeline. FACS, fluorescence activated cell sorting. **(B)** Diagram of the strategy for identifying genotyping markers on the reference genome by mapping short reads and markers in gametes by mapping scRNA-seq reads across a large number of gametes to the reference genome. GMRs, genotyping markers on reference genome. **(C)** Diagram of the identification of potential CO events after the alignment of the scRNA reads from each gamete to the phased reference genome. **(D)** An example of genotype definition by markers in a real pollen nucleus, e.g., cell barcode AAGACTCTCATCCTAT.

### Single-cell RNA sequencing of pollen allows the high-throughput identification of genome-wide COs

We identified genome-wide CO events by conducting 10X Genomics single-cell RNA sequencing (scRNA-seq) on the nuclei extracted from pollen grains of *R. breviuscula* (**Figure S4, see Methods**). After pre-processing, we obtained individual sequence data for 4,392 *R. breviuscula* pollen nuclei. After removing residual doublets and cells with low number of reads (**Figure S4, see Methods**), we obtained a final set of 1,641 pollen nuclei with at least 400 markers (∼1 marker/Mb). These markers (median resolution ∼1 marker/542 kb) covered almost the entire length of all five chromosomes (**Figure S5B**), ensuring genome-wide CO detection. We detected 4,047 COs in the 1,641 pollen nuclei by inspecting genotype conversions, as indicated in **Figure 4C,D** (**Figure S6**). Overall, we delineated a complete and detailed pipeline to detect COs in an economical way by high-throughput scRNA-seq of gametes from a single heterozygous individual (**Figure 4**).

### Repeat-based holocentromeres show a U-shape recombination landscape

Counting the occurrence of COs in chromosome-wide genomic intervals across all pollen nuclei, we computed the CO rates along chromosomes. This is the first recombination map for *R. breviuscula*, and the first in any species with known repeat-based holocentromeres (**Figure 5A**). The overall resolution of location interval between COs was ∼1.5-Mb (median) and ∼2.24-Mb (mean). The landscape contained large regions or high- and low-recombination frequencies, i.e., recombination domains. Most regions with high recombination rates were located at distal chromosomal regions, while central chromosomal regions showed lower recombination rates. Unexpectedly, the recombination landscape of holocentric *R. breviuscula* resembled a U-shape distribution of COs, which is commonly present in monocentric models (**Figure 5A,B**). Remarkably, chromosomes 1 and 2 each had only one high-CO domain at one chromosomal end, while the other end showed a lower CO level similar to the central region. The other three chromosomes harboured two high-recombination domains at both ends (**Figure 5A**). We thus reveal an uneven distribution of CO rates (at chromosomal scale), despite the uniform distribution of centromeric units (see below; Hofstatter et al. ^32^). The total linkage map length was 246 cM, corresponding to ∼50 cM per chromosome (**Figure 5B**).

**Figure 5.**
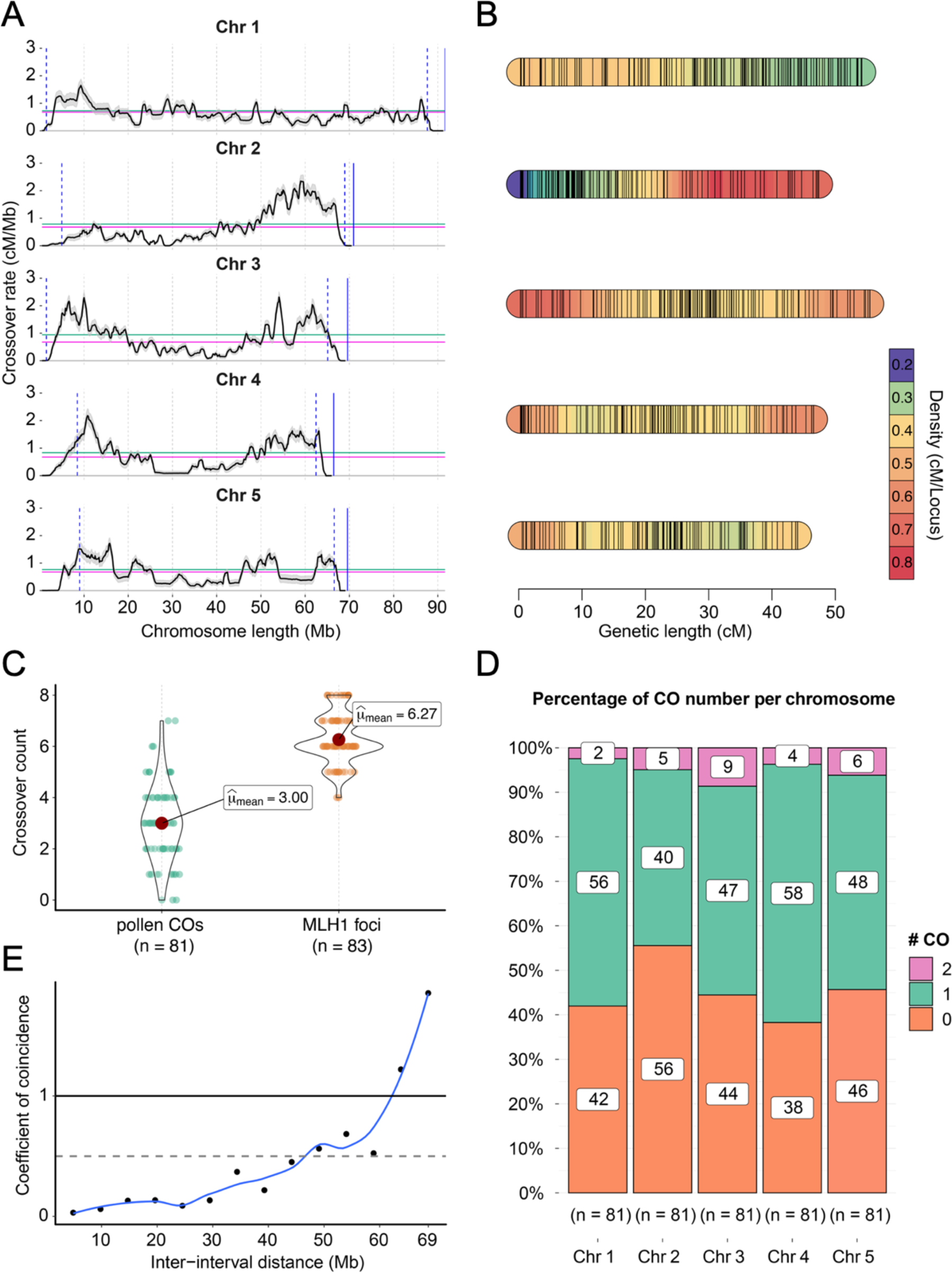
Meiotic recombination dynamics in *R. breviuscula* derived from single-pollen sequencing. **(A)** The first recombination landscape of the five chromosomes in *R. breviuscula* achieved by computing COs from 1,641 pollen nuclei. Black line displays the CO rate, which is the mean of 500 random samplings for each CO interval. Shadow ribbons indicate one standard deviation from mean CO rates. Blue dashed vertical lines: start and end of confident CO rate computation (**Figure S7**). Blue solid vertical line: chromosomal end. Magenta horizontal line: genome-wide average CO rate. Green horizontal line: chromosome-wide average CO rate. **(B)** Genetic linkage map with genetic length density indicated by colouring. A set of 705 markers was selected using a 500-kb sliding window through all markers defined against the reference (**see Methods**). **(C)** CO number derived by counting CO events from the bioinformatic analysis and the number of MLH1 foci from cytological observations. **(D)** Distribution of CO number for each chromosome. Note the higher incidence of double COs on chromosome 3. **(E)** CoC curve in pollen nuclei (*n* = 1,641). Chromosomes were divided into 15 intervals, with random sampling at CO intervals, to calculate the mean coefficient of coincidence of each pair of intervals.

We compared CO numbers estimated from DNA sequencing and the number of MLH1 foci observed by cytology. To have a precise estimation of CO number, we counted only those COs from pollen nuclei with more than 2,000 markers (*n* = 81). On average, we detected around three COs per haploid gamete, or 0.6 CO per chromatid (**Figure 5C,D**). As gametes only have one chromatid from each recombined chromosome, the number of pollen-detected COs should be approximately half of COs that occur in the meiocytes. Interestingly, we found an approximately similar number of MLH1 foci and COs detected in our genetic analysis (**Figure 5C**), suggesting that most of the COs formed in *R. breviuscula* are of class I. Furthermore, all chromosomes had exactly one CO in half of these gametes (*n* = 81), while double COs appeared in only 5% of the 81 gametes considered (**Figure 5D**). Chromosome 3 showed the highest frequency of double COs (9%; **Figure 5D**), which conferred it the longest genetic length among all *R. breviuscula* chromosomes (55 cM; **Figure 5B**). This is especially remarkable considering that chromosome 1 (53 cM) is physically longer than chromosome 3 (by 20 Mb).

We also tested whether CO interference occurred in *R. breviuscula*. We used a Chi-square goodness-of-fit test to investigate whether the CO number on each chromosome follows a Poisson distribution, which revealed a significant discrepancy between observed and expected CO numbers (**Figure S8A**). This result shows that COs are not randomly distributed but under-dispersed, based on the negative alpha values from dispersion tests, that could be the effect of CO interference. We also computed the coefficient of coincidence (CoC) of COs across the genome, which measures the observed frequency of double COs over their expected frequency (**see Methods**). The CoC curve of all chromosomes showed that the coefficients are below 1 for genomic intervals with distances less than around 60 Mb (**Figure 5E; Figure S8B**), showing that the frequency of double COs is lower than expected. This result indicates the presence of strong CO interference in *R. breviuscula*.

### Broad-scale recombination landscape is independent of holocentromere distribution and (epi)genomic features

We compared the broad-scale recombination landscape with all known (epi)genetic features to determine whether any specific feature would explain the CO distribution of *R. breviuscula*. A chromosome-wide comparison of the recombination landscape of *R. breviuscula* revealed no apparent correlation with the uniform holocentromere distribution and other genomic (genes, TEs, SNP densities, or with GC content) and epigenomic features (such as H3K4me3, H3K27me3, H3K9me2 or DNA methylation). To estimate whether fast-evolving genes correlated with the regions showing higher recombination frequencies, we also compared the Ka/Ks ratio (measurement of the relative rates of synonymous and nonsynonymous substitutions at each gene). Notably, the Ka/Ks ratio across the chromosomes was rather uniform, and we did not find any bias towards the chromosome ends (**Figure S9**). No (epi)genomic feature showed strong correlation with CO distribution, as they are all found uniformly distributed along *R. breviuscula* chromosomes (**Figure 6A,B**). These results indicate that, at broad scale, meiotic recombination occurs independently of chromosome-wide holocentromere distribution and (epi)genomic features.

**Figure 6.**
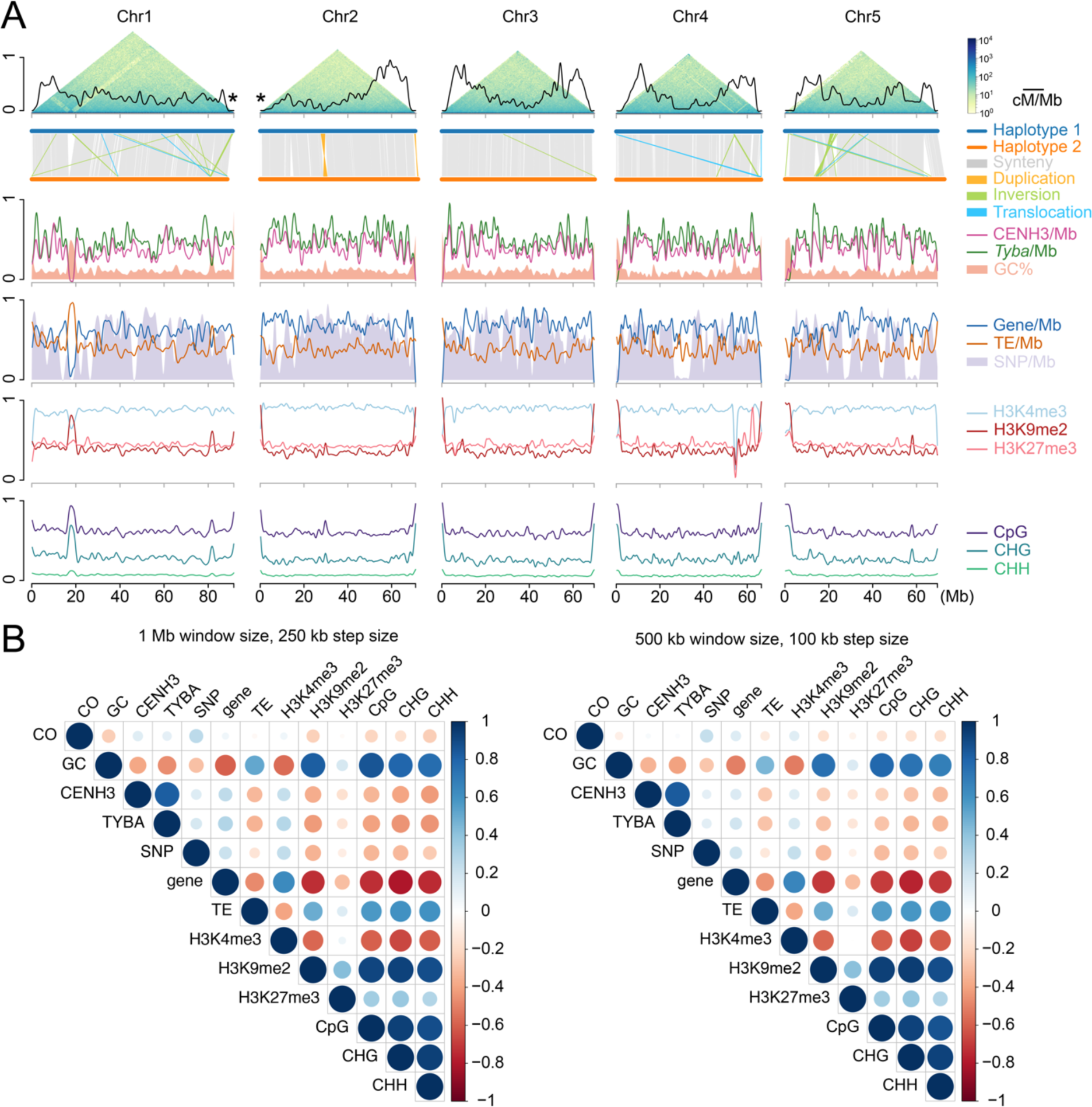
Broad-scale correlation of the recombination landscape and (epi)genetic features in *R. breviuscula*. **(A)** Chromosome distribution of the CO rate coupled with different (epi)genetic features. Top: recombination landscape (black line) created with sliding windows of 500 kb at a step of 50 kb, with COs detected in all single-pollen nuclei (*n* = 1,641), coupled with Omni-C chromosome conformation capture contacts. The terminal locations of the *35S rDNA* loci on chromosomes 1 and 2 are indicated by asterisks. For the x-axes, the coordinates were based on the haploid 1 assembly *R. breviuscula*. For the y-axes, all features were scaled [0,1], with 1 indicating a maximum of 2.34 for recombination frequency (cM/Mb), 5 for *Tyba* density, 6 for CENH3 density, 7205 for SNP density, 88 for gene density, and 227 for TE density. GC [33.3, 46.6], H3K4me3 [–1.494, 0.231], H3K9me2 [–1.20, 1.84], and H3K27me3 [–0.671, 0.491] are scaled to [0,1] by their minima and maxima. mCG, mCHG and CHH are original values (0 to 100%) and are scaled so 1 represents 100%. COs are almost completely absent in a large inversion in chr2:30–35 Mb, while in homozygous regions we could not confidently call COs, for example in chr4:25–35 Mb. The large variants were confirmed within Hi-C contact maps (Figure 3C). Asterisks at the chromosome ends of chromosomes 1 and 2 indicate the position of the *35S rDNA* clusters in the assembly and confirmed by FISH (**Figure S13**). **(B)** Correlation matrix illustrating 4,047 COs correlation with all available (epi)genetic features. Positive correlations are displayed in blue and negative correlations in red. Colour intensity and the size of the circle are proportional to the correlation coefficients. In the right side of the correlogram, the legend colour shows the correlation coefficients and the corresponding colours. Pearson correlation coefficients for each pair of all features under 1-Mb smoothing window and 250-kb step size (left) and 500-kb smoothing window and 100-kb step size (right): specifically, mean CO rates, mean GC contents, CENH3 peak density, *Tyba* array density, SNP density, TE density, H3K4me3 RPKM, H3K9me2 RPKM, H3K27me3 RPKM, mean CpG, mean CHG, and mean CHH.

### Absence of centromere effect sheds light on the fine-scale epigenetic CO regulation

As we did not find any correlation between the CO distribution and (epi)genetic features at a broad genomic scale, we tested for local centromere effects on CO designation in *R. breviuscula*. Although our scRNA-seq strategy is useful for delineating the recombination landscape and CO dynamics, the overall CO resolution obtained was low (median size of the location interval ∼1.5 Mb, mean ∼2.24 Mb), which does not allow for a precise analysis of a potential centromere effect in this particular case. To achieve precise CO resolution, we performed manual self-pollination in *R. breviuscula*. Due to its high self-incompatibility, we obtained only 63 F_1_ plants by selfing; we sequenced these to 3x coverage, which allowed us to detect 378 CO events at a very high resolution (median 334 bp, mean ∼ 2 kb). Overall, we obtained results of COs number and distribution similar to our single-pollen sequencing strategy, confirming the robustness of our analysis (**Figure S10**). We observed an increase in the genetic map length in the F_1_ offspring, suggesting that heterochiasmy occurs in *R. breviuscula* and that female meiosis might have slightly higher CO frequencies than male meiosis (**Figure S10A,B**). We estimated the average CO number to be 6 in the F_1_ offspring, exactly double the average number estimated from single-pollen nuclei data (**Figure S10C,D**).

The holocentromeres in *R. breviuscula* are repeat-based, i.e., each centromeric unit is based on a specific array of the holocentromeric repeat *Tyba* associated with CENH3, with average sizes of ∼20 kb and average spacings of ∼400 kb, where each chromosome harbours hundreds of individual centromeric units (**Figure 7A,B**). Remarkably, we found the same epigenetic centromere identity in *R. breviuscula* (**Figure 7C**) as reported for *R. pubera*^32^. This organisation makes it possible to identify centromeric units at the DNA level by annotating *Tyba* repeat arrays (**Figure 7B**). We then computed the observed *versus* expected by random distribution fine-scale CO position across all available chromatin marks and genetic features. We found that COs are more frequently formed at H3K4me3 peaks and genes than what expected by random distribution (**Figure 7D; Figure S11**). Within genic regions COs were preferentially formed at the promoter regions (**Figure 7E**). Remarkably, COs were mostly suppressed at core centromeric units and heterochromatic regions (**Figure 7D,F; Figure S11**). However, after computing the distances between the CO break intervals and the corresponding nearest *Tyba* arrays/CENH3 domains, the COs did not show a tendency to be positioned away from or close to centromeric units (**Figure 7G**), indicating the absence of a centromere effect and that the proximity to a centromeric unit does not affect CO formation. Moreover, we found five cases of a CO being placed inside a region containing reduced *Tyba* repeats and CENH3-positive chromatin (**Figure 7H**). Our results point to the exciting finding that local CO formation in *R. breviuscula* is abolished at repeat-based centromeric units but enriched at genic promoter regions, supporting the role of chromatin features at fine scale in contrast to the absence of correlation at broad scale (**Figure 7I**).

**Figure 7.**
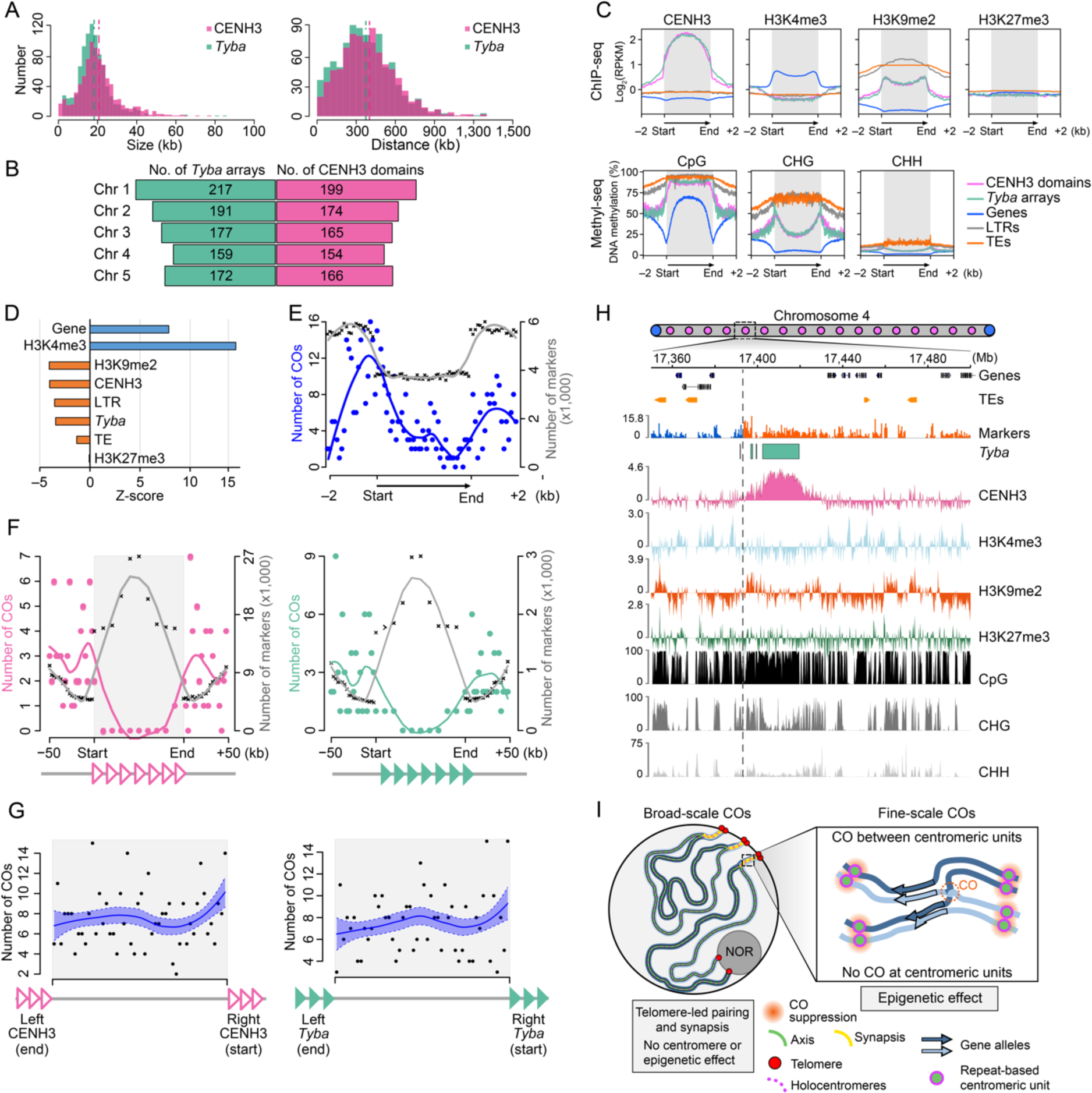
Epigenetic regulation and fine-scale correlation of CO positions in repeat-based holocentromeres. **(A)** Size (left) and spacing (right) length distribution of CENH3 domains and *Tyba* arrays. CENH3 domain median size is 19156 bp and the mean size is 20697 bp. The median of *Tyba* array size is 17424 bp and the mean is 18220 bp. CENH3 domain median spacing is 378,467 bp and the mean is 401,763 bp. The median of *Tyba* array spacing is 354,850 bp and the mean is 374,310 bp. **(B)** Number of *Tyba* arrays (left) and CENH3 domains (right) for each chromosome annotated in the reference haplotype genome. **(C)** Enrichment of CENH3, H3K4me3, H3K9me2, and DNA methylation in the CpG, CHG, and CHH contexts from the start and end of different types of sequences: CENH3 domains (magenta), *Tyba* repeats (green), genes (grey line), LTRs (yellow-green), and TEs (orange). ChIP-seq signals are shown as log2 (normalised RPKM ChIP/input). Grey boxes highlight the modification enrichment over the body of each sequence type. **(D)** Z-score of the overlapped CO numbers with different (epi)genetic features to the 5,000 simulations of randomly distributed COs. Positive z-score indicates that COs overlap with H3K4me3 and genes more frequently than expected under the hypothesis of random distributed COs along chromosomes. Negative z-score implies the contrary. The higher the absolute z-score, the more deviation is observed. **(E)** Within genic regions, CO frequency (blue line) is higher in promoter regions or after the transcription termination site (TTS), but lower in gene bodies, independent of marker density (grey line). TSS, transcription start site. **(F)** Within CENH3 domains (left) and *Tyba* arrays (right) CO frequency is remarkably suppressed, despite relative high marker density. **(G)** Random distribution of the relative distance of CO positions to the end of the left and to the start of the right CENH3 domain (left) or *Tyba* array (right). The median of CO resolution is 334 bp and the mean is about 2 kb. Correlation analysis performed using data from 63 F1 recombinant offspring and a total of 378 COs. Pink-bordered and green-filled triangles represent CENH3 domains (pink) and *Tyba* repeat arrays (green), respectively. **(H)** Magnified view of one of the five COs placed within a region containing CENH3-positive chromatin and *Tyba* repeats. CO resolution in this case 200 bp. CO is indicated by the grey dashed line showing the haplotype switch (blue to orange) in the Marker density track. **(I)** Model for CO formation at (left) broad- and (right) fine-scale. Telomere-led synapsis leads to an early loading of HEI10 at chromosomal ends that can potentially favour COs at distal regions, while 35S-rDNA harbouring chromosome ends do not show early synapsis and thus have less probability of making COs. At local scale, COs are suppressed at core centromeric units, but not at their vicinity, where COs can be placed anywhere between two centromeric units. Remarkably, COs were preferentially placed at gene promoter regions.

### Spatiotemporal dynamics of chromosome pairing and synapsis explains the broad-scale recombination landscape

We hypothesised that pairing and synapsis progression might contribute to the U-shaped recombination landscape observed in *R. breviuscula*. To investigate this question, we performed immunolocalisation with antibodies against ZYP1, ASY1, and HEI10 and fluorescence *in situ* hybridisation (FISH) for telomeres on meiocytes. Signals detected for ZYP1, ASY1 and telomere probes indicated a tendency for telomeric signals to cluster together in one location, forming the typical “bouquet” ^52, 53^. Near this structure, we observed ZYP1 signal, representing synapsed chromosomes elongating from telomeres until they reach the area of the nucleolus that is not yet synapsed. Here, the linear signal of ASY1 was still present and represented unpaired chromosomes (**Figure 8A; Figure S12**). When using telomeric probes and antibodies against ZYP1 and HEI10, we determined that the first synapsed regions (ZYP1-stained) were also first loaded with HEI10 in the proximity of chromosome ends, exhibiting a high-intensity linear signal (**Figure 8B,C**). We consistently observed a few telomeres that did not participate in the bouquet, coming from the terminal ends of chromosomes 1 and 2 that harbour the *35S rDNA* loci; instead, these chromosome ends localised in the nucleolus (**Figure 6A; Figure S13**). Remarkably, the nucleolus-positioned telomeres showed delayed ZYP1 loading – if it happens at all – compared to the telomeres involved in the bouquet (**Figure 8B,D**). Thus, the broad-scale recombination landscape in *R. breviuscula* is better explained by early synapsis and HEI10 loading on the terminal regions of early synapsed chromosomes rather than by any association with a centromere effect or (epi)genetic features (**Figure 7I**). A similar spatiotemporal asymmetry of synapsis has been recently proposed to explain the distal bias of meiotic COs in wheat^54^.

**Figure 8.**
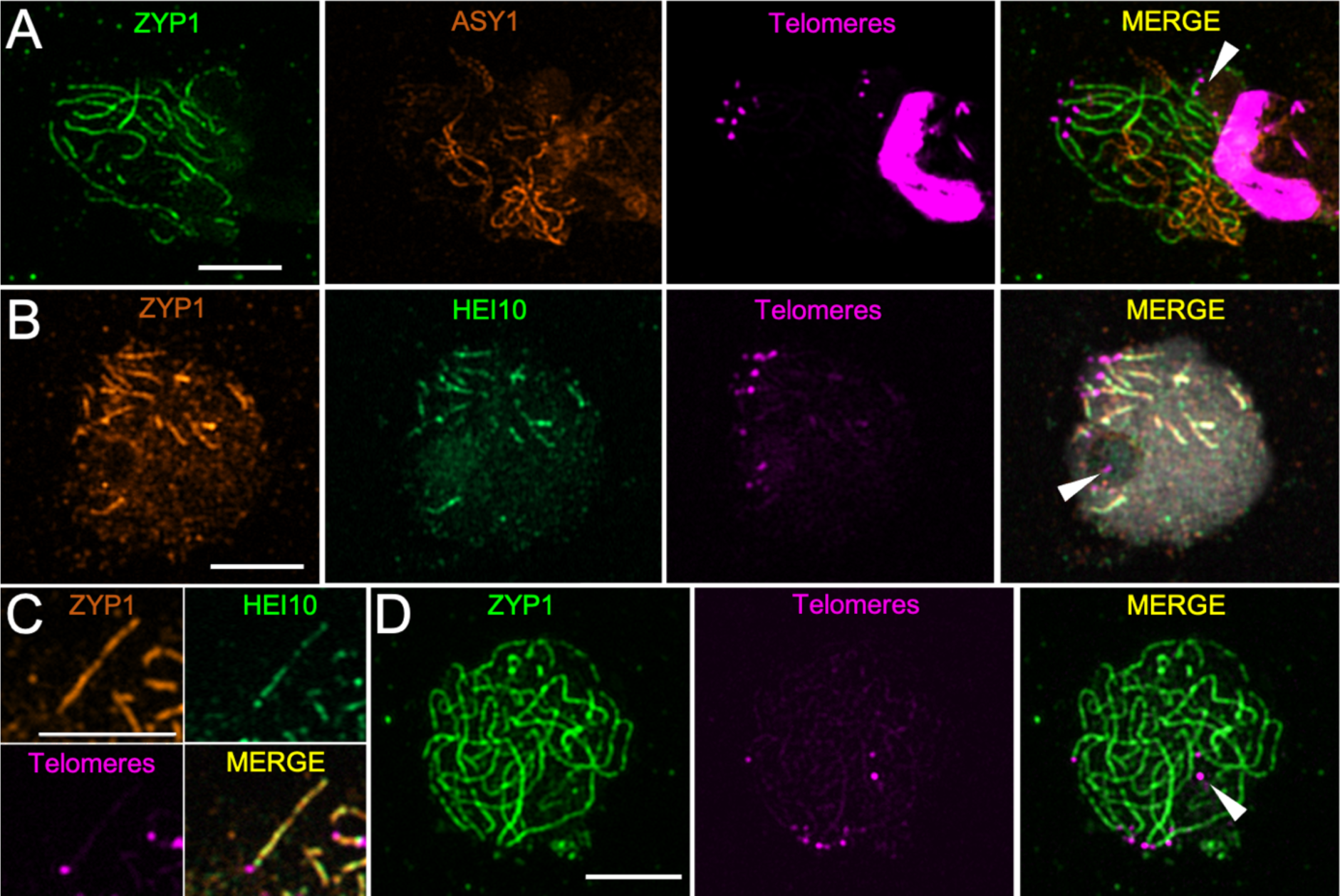
Immunolocalisation of ZYP1, ASY1, HEI10 and telomere-FISH. **(A)** Telomeres (magenta) cluster in a bouquet (white arrowhead) on one side of the cell, where ZYP1 (green) elongating as the SC is being assembled. ASY1 (orange) represents unpaired chromosomes not yet reached by ZYP1. **(B)** As ZYP1 (orange) lines elongate from the telomeres (magenta), HEI10 (green) is quickly loaded onto paired chromosomes, while some telomeres localise to the nucleolus (white arrowhead) and lack the ZYP1 and HEI10 signals. **(C)** Detail of synapsis progression: As soon as the SC (orange) is assembled, HEI10 (green) is loaded. **(D)** In late pachytene, ZYP1 (green) occupies the whole chromosomal length, and telomeres (magenta) are still clustered in the bouquet or at the nucleolus (white arrowhead). Scale bars, 5 µm.

## Discussion

Deciphering the mechanisms controlling CO formation and distribution is key to understanding one of the main driving forces for genetic diversity in eukaryotes: meiotic recombination. By combining comprehensive immunocytochemistry, chromatin, and genetic analyses of the recombination dynamics in the holocentric plant *R. breviuscula*, we determined that telomere-led pairing and synapsis can explain the U-shaped recombination landscape observed. This result is consistent with the bouquet formation reported in many organisms, where synapsis and DNA double-strand breaks (DSBs) required for COs are mostly initiated from the telomeres^52, 53^. Such telomere-led mechanisms have already been proposed to influence the location of COs to be more likely at the chromosome ends than the centres (see Haenel et al. ^36^ and references therein). Considering the marked conservation for bouquet formation and synapsis progression in *R. breviuscula*, and the position of high- and low-recombination domains, we propose that pairing itself, and possibly the observed telomere-led HEI10 loading dynamics (**Figures 2 and 8**), are the driving force that shapes the recombination landscape in this species. This early loading at ends might create a bias that increases CO rates at the distal regions of the chromosomes, whether a centromere is present or not. Recently, a “coarsening” model for the behaviour of HEI10 has been proposed. In this model, enhanced loading of HEI10 at the chromosome ends leads to increased COs. As the amount of loaded HEI10 accounts for the increased coarsening over time, early loading at the chromosome ends would accelerate the maturation of recombination intermediates compared to the interstitial regions of the chromosomes^46, 47^.

Moreover, we observed a gradual reduction in CO rates from the regions directly adjacent to telomeres in *R. breviuscula*. Similar to the centromere effect, a telomere effect is proposed to commonly occur across eukaryotes^17, 36^ and might be explained by the coarsening model. We hypothesise that, as pairing and synapsis progress and finally involves the whole length of the chromosomes, recombination intermediates are affected by the coarsening coming from both ends. Eventual recombination intermediates at the telomeres will therefore be less subject to the effect of the coarsening compared to more internal COs. Additionally, the phenomenon of CO interference lowers the recombination frequencies at the centre of chromosomes, with the distal regions having already been designated for COs. The model described here would explain the behaviour of chromosomes 3, 4 and 5; however, the *35S rDNA*-harbouring distal regions of chromosomes 1 and 2 do not participate in bouquet formation, as they stay at the nucleolus. Remarkably, these two chromosomal ends are also characterised by the lowest recombination frequencies. For the model plant *A. thaliana*, it has been proposed that ribosomal DNA is not involved in synapsis and recombination and that these regions localise in the nucleolus^55, 56^. Indeed, we observed that these telomeres situated in the nucleolus were involved later in synapsis than those that cluster in the bouquet. This late involvement in synapsis means a potential delay in DSB formation and HEI10 loading, which is consistent with the lower recombination frequency observed at the *35S rDNA*-harbouring ends of chromosomes 1 and 2. Here, we show that the CO distal bias is present even in the absence of compartmentalised chromosomal features, strongly suggesting that telomere-led pairing and synapsis initiation alone can impose CO bias^54, 57–59^. However, we cannot exclude that other factor, like a different density of DSBs along chromosomes might contribute to the U-shape distribution of COs^60^. Future experiments in organisms with repeat-based holocentromeres will be important to identify conserved and adapted mechanisms about the role of centromeres in the spatiotemporal dynamics of meiotic DSB formation and HEI10 loading.

In the new era of highly-accurate long read genomics, haplotype-phased genomes are routinely available. By applying high-throughput single-cell RNA sequencing to individual pollen nuclei, we provide a powerful pipeline that can be used to investigate CO frequencies in any available gamete of any heterozygous individual with an available phased genome. Using haplotype-specific markers, we detected and mapped CO events from thousands of gametes for the first time in a species with repeat-based holocentromeres. Unexpectedly, the recombination rates were not homogeneously distributed along the chromosomes of *R. breviuscula*, as one might expect from its absence of chromatin compartmentalisation and the uniform distribution of (epi)genetic features (this study; ^32^). Instead, we observed regions of higher recombination frequencies (recombination domains) mainly at distal chromosomal regions, similar to the observed in most eukaryotes, including the holocentric *C. elegans*^35, 36, 61^. A recent study showed that the megabase-scale CO landscape in *A. thaliana* is mostly explained by association with (epi)genetic marks beyond a centromere effect, with open chromatin states showing the highest positive correlation with CO formation^15^. While single nucleotide polymorphisms only showed a rather local effect on CO positioning^15, 62^. In contrast to these organisms, we could only find correlation of CO positioning with centromere and (epi)genetic features at a very fine-scale.

At local scale, COs preferentially formed within gene promoter regions rather than in neighbouring transcribed gene bodies, TEs and centromeres. This result appears to hold true for several eukaryotes and may be related to open chromatin states^14, 15, 63^, suggesting that CO regulation at fine-scale is associated with similar (epi)genetic factors independent of the chromosome organisation. In contrast, the absence centromere effect found in *R. breviuscula*, which seems to suppress CO formation only at the core centromeric units but not at their vicinity, is likely due to the closed chromatin state of centromeric chromatin as marked by high DNA methylation, as also found within TEs. Our findings suggest that the pericentromeric inhibition of COs observed in many eukaryotes^17, 64^ is likely a secondary effect of pericentromeres evolution and not a direct effect of centromeric chromatin. We show that by using a holocentric species, where the lack of localized centromeres and compartmentalised chromosome organisation, features that can potentially mask the factors underlying CO patterning, can reveal important insights into CO control mechanisms. Understanding the molecular mechanisms of CO control in holocentric organisms will potentially unveil new strategies to address meiotic recombination within centromere proximal regions of monocentric chromosomes that rarely recombine.

## Data and code availability

All sequencing data used in this study have been deposited at NCBI under the Bioproject no. XXXXXXX and are publicly available as of the date of publication. The reference genomes, sequencing data, annotations and all tracks presented in this work are made available at DRYAD LINK. All other data needed to evaluate the conclusions in the paper are provided in the paper and/or the supplemental information. The original code for the construction of recombination maps from single-cell RNA sequencing is available at https://github.com/Raina-M/detectCO_by_scRNAseq. Any additional information required to reanalyse the data reported in this paper is available from the corresponding author upon request.

## Acknowledgements

We thank Raphaël Mercier for the insightful comments and critical reviewing on the manuscript. We thank Neysan Donnelly for reviewing the manuscript. This study was financially supported by the Max Planck Society (core funding to AM) and the Deutsche Forschungsgemeinschaft (DFG, grant number MA 9363/2-1 to AM). MZ is financially supported by the DFG (grant number MA 9363/2-1). We thank the PhD fellowship awarded to G.T. from the DAAD/India. YMS received financial support from PROBRAL (CAPES/DAAD) program (grant number 88881.144086/2017-01). JAC is financially supported by the Marie Skłodowska-Curie Individual Fellowship PrunMut (grant number 789673).

## Author contributions

AM conceived the research program and coordinated the analyses. MC performed all cytogenetic analyses and microscopy. MC isolated the pollen nuclei and generated sequencing libraries with assistance from JAC. MZ performed all single-cell RNA sequencing and recombination-related analyses with assistance from HS. GT performed the ChIP-seq analysis. YMS performed the immuno-FISH analysis. TL and KFXM performed the gene annotation and Ka/Ks ratio analysis. MM operated the FACS machine. BH performed all sequencing. KS supervised the single-cell analysis. MC, MZ and AM wrote the first manuscript draft with input from all authors. All authors approved the final version of the manuscript.

## Methods

### DNA isolation from pollen nuclei, 10X Genomics scRNA-seq library preparation and sequencing

Protocols were adapted from Campoy et al. (2020). Briefly, to release pollen grains, anthers from fully developed flowers of *R. breviuscula* and *R. tenuis* (for multiplex purposes) were harvested and submerged in woody pollen buffer (WPB; Loureiro *et al.* 2007). The nuclei were extracted using a modified bursting method. The solution containing the pollen grains was pre-filtered with a 100-µm strainer, and the pollen was crushed on a 30-µm strainer (Celltrics). The isolated nuclei were gathered in WPB and stained with DAPI (1 µg/ml) before being sorted using a BD FACSAria Fusion sorter with a 70-µm nozzle and 483-kPa sheath pressure. A total of 10,000 nuclei were sorted into 23 µl of sheath fluid solution and loaded into a 10X Chromium controller, according to the manufacturer’s instructions. A library was created according to the chromium single-cell 3′ protocol. A CG000183 Rev A kit from 10X Genomics was used for library preparation. The library was sequenced (100 Gb) on an Illumina NOVAseq instrument in 150-bp paired-end mode.

### Whole-genome sequencing (WGS) of F_1_ recombinant offspring

To obtain a recombinant population of *R. breviuscula* plants, young inflorescences of the heterozygous reference *R. breviuscula* were bagged to force self-pollination. Due to its high self-incompatibility, only 63 F_1_ plants were obtained, and they were sequenced to 3X coverage (∼2 Gb) using an Illumina NextSeq2000 instrument in 150-bp paired-end mode.

### Anther fixation and immunocytochemistry

Immunostaining was performed as described by Cabral et al. (2014), with some modifications. Anthers of *R. breviuscula* were harvested and fixed in ice-cold 4% (w/v) paraformaldehyde in phosphate buffered saline (pH 7.5; 1.3 M NaCl, 70 mM Na_2_HPO_4_, 30 mM NaH_2_PO_4_) for 90 min. The anthers were separated according to their size and were dissected to release the meiocytes onto glass slides. The meiocytes were squashed with a coverslip that was later removed using liquid nitrogen. The slides were stained with mounting solution (Vectashield + 0.2 µg DAPI) to select the meiotic stages of interest, after which they were blocked with a 1 h incubation in 3% (w/v) bovine serum albumin in PBS + 0.1% (v/v) Triton X-100 at 37°C. The antibodies used were anti-AtASY1 raised in rabbits (inventory code PAK006)^40^, anti-AtMLH1 raised in rabbits (PAK017)^66^ and anti-RpCENH3 raised in rabbits^30^. The anti-ZYP1 was raised in chickens against the peptide EGSLNPYADDPYAFD of the C-terminal end of AtZYP1a/b (gene ID: At1g22260/At1g22275) and affinity-purified (Eurogentec) (PAK048). The anti-RpREC8 was a combination of two antibodies raised in rabbits against the peptides CEEPYGEIQISKGPNM and CYNPDDSVERMRDDPG (gene ID: RP1G00316120/RP2G00915110/RP4G01319620/RP5G01638170) and affinity-purified (Eurogentec). The anti-RpHEI10 was a combination of two antibodies raised in rabbits against the peptides CNRPNQSRARTNMFQL and CPVRQRNNKSMVSGGP (gene ID: RP3G01271190/RP3G01008630/RP1G00269340/RP2G00699130) and affinity-purified (Eurogentec). Each primary antibody was diluted 1:200 in blocking solution. The slide-mounted samples were incubated with the primary antibodies overnight at 4°C, after which they were washed three times for 10 min with PBS + 0.1% (v/v) Triton X-100. The slides were incubated with the secondary antibodies for 2 h at room temperature. The secondary antibodies were conjugated with Abberior STAR ORANGE or Abberior STAR RED (1:250; Abberior) before being washed again three times for 10 min with PBS + 0.1% (v/v) Triton X-100 and allowed to dry. The samples were prepared with 10 µl of mounting solution (Vectashield + 0.2 µg DAPI), covered with a coverslip, and sealed with nail polish for storage. Images were taken with a Zeiss Axio Imager Z2 with Apotome system for optical sectioning or with a Leica Microsystems Thunder Imager dMi8 with Computational Clearing. The images were deconvolved and processed with Zen 3.2 or LAS X software.

### Sequential immunostaining and fluorescence *in situ* hybridisation

Immuno-FISH was performed following Baez et al. (2020). The best slides obtained from immunostaining, as described above, were selected for FISH using a telomeric probe. The slides were washed with 1× PBS for 15 min, postfixed in 4% (w/v) paraformaldehyde in PBS for 10 min, dried with 70% (v/v) and 100% ethanol for 5 min each and probed with direct-labelled telomeric sequence (Cy3-[TTTAGGG]_5_; MilliporeSigma). The hybridisation mixture contained formamide (50% w/v), dextran sulphate (10%, w/v), 2× SSC and 50 ng/μl of telomeric probe. The slides were denatured at 75°C for 5 min. Stringency washes were performed following^68^ to give a final stringency of approximately 72%. The slides were counterstained with 10 µl of mounting solution (Vectashield + 0.2 µg DAPI), and images were captured as described above.

Mitotic and meiotic chromosome spreads were performed as described by Ruban et al. (2014), with some modifications. Briefly, tissue samples were fixed in 3:1 (ethanol:acetic acid, v/v) solution for 2 h with gentle shaking. The samples were washed with water twice for 5 min and treated with an enzyme mixture (0.7% [w/v] cellulase R10, 0.7% [w/v] cellulase R10, 1.0% [w/v] pectolyase, and 1.0% [w/v] cytohelicase in citric buffer) for 30 min at 37°C. The material was immersed in freshly prepared 60% (v/v) acetic acid, and the samples were dissected on slides under a binocular microscope. The slides were placed on a hot plate at 50°C and the samples were spread by hovering a needle over the drop of acetic acid without touching the slide. After spreading the cells, the fixation was completed by dropping fresh 3:1 (v/v) fixative on the slides and immersing them in 60% (v/v) acetic acid for 10 min. The slides were dehydrated in 100% ethanol and air-dried, ready for future applications.

### Haplotype phasing and scaffolding

A phased chromosome-level genome of *R. breviuscula* was assembled using PacBio HiFi and Hi-C data available from Hofstatter *et al.* (2022) under NCBI Bioproject no. PRJNA784789. First, a phased primary assembly was obtained by running Hifiasm^50^ using as inputs the 30 Gb of PacBio HiFi reads (∼35X coverage per haplotype) in combination with Dovetail Omni-C reads, using the following command: hifiasm -o Rbrevi.phased.asm.hic --h1 hic.R1.fastq.gz --h2 hic.R2.fastq.gz hifi.reads.fastq.gz. The phased assemblies of each individual haplotype were further scaffolded to chromosome scale using Salsa2^70^, followed by successive rounds of manual curation and re-scaffolding. The genome sizes of haplotypes 1 and 2 were 418,624,405 and 390,890,712 bp, respectively. Both haplotypes comprise five chromosomes with a length of ∼370 Mb in total, as well as other unplaced sequences (**Table S1**).

### Definition of allelic SNPs as genotyping markers on the phased reference genome

To define genotyping markers for *R. breviuscula*, all available (NCBI Bioproject no. PRJNA784789) raw Illumina HiSeq3000 150-bp paired-end reads (25,899,503,075 bases, ∼54X coverage) were first mapped to the five pseudochromosome scaffolds in haplotype 1 of the phased reference genome using bowtie2 (v2.4.4)^71^. The alignment file was further sorted with SAMtools (v1.9)^72^. The alignments of short reads to the reference genome were used for SNP calling by ‘bcftools mpileup’ and ‘bcftools call’ (v1.9)^72^ (with the --keep-alts, --variants-only, and --multiallelic-caller flags enabled). A total of 1,404,927 SNPs excluding indels were derived. To distinguish the two haplotypes using these SNPs, only allelic SNPs were selected as markers for genotyping; therefore, variant information was collected, including mapping quality, alternative base coverage, and allele frequency resulting from SHOREmap conversion (v3.6)^73^, which converts SNP files (.vcf) into a read-friendly, tab-delimited text file. A final set of 820,601 alleles fulfilling certain thresholds (mapping quality > 50; 5 ≤ alternative base coverage ≤ 30, 0.4 ≤ allele frequency ≤ 0.6) was selected as markers (**Figure 4B; Figure S3**).

### Pre-processing single-cell DNA sequencing data from pollen nuclei

Raw scRNA-seq data usually include barcode errors and contaminants such as doublets and ambient RNA. In the present study, cell barcodes (CBs) were first corrected in these data using ‘bcctools correct’ (v0.0.1) based on 10X v3 library complete barcode list with options “--alts 16 --spacer 12” because of the 16-bp CB and 12-bp unique molecular identifier (UMI). After correction, 952,535 viable CBs were detected. This step also truncated the CBs and UMIs from every pair of scRNA-seq reads. After counting the occurrence of CBs, the number of read pairs under each CB was determined. To ensure a sufficient number of reads for SNP calling, only CBs appearing more than 5,000 times were used for the subsequent analyses. Finally, each CB was seen as one viable cell, and reads corresponding to the CB were assigned to this cell (demultiplexing). A total of 8,001 viable cells were ultimately identified, with 365,771,748 (77.25% of all raw scRNA-seq) read pairs included.

### Alignments of single-pollen DNA sequences to genome and deduplication

To identify genotyping markers in the *R. breviuscula* gametes, scRNA reads of the pollen nuclei were first mapped to the haplotype 1 chromosomes (**Figure 4B**) using hisat2 (v2.1.0)^74^. Specifically, each cell-specific pair of reads was merged as one single-end FASTQ file, and hisat2 was run under single-end mode (-U) because the SNP-calling approach does not detect SNPs on reads whose mated reads are not mapped. Before further analyses of the alignment results, UMIs were previously extracted from the read alongside the CBs; hence, a fast UMI deduplication tool, UMIcollapse^75^, was employed to remove the PCR duplicates by collapsing reads with the same UMIs.

The sequencing library was prepared for mixed pollen nuclei of *R. breviuscula* and *R. tenuis* to enable multiple-potential analyses. The addition of gametes from *R. tenuis* was done for multiplexing purposes, and they will be analysed in another study. To discriminate the single-cell data between the two species, we used a straightforward approach without gene expression profiling: For each cell, a) the DNA sequences were mapped to both the *R. breviuscula* and *R. tenuis* chromosomal genomes; and b) the alignment rates between the two species were compared to decide the cell identity (**Figure S4**). The alignment rates to *R. breviuscula* and *R. tenuis* were both bimodal distributions (**Figure S4A,B**); therefore, these cells can be grouped solely based on their mapping rates. It was estimated that 4,733 cells were from *R. breviuscula* and 2,709 cells were from *R. tenuis* (**Figure S4C**) based on the alignment fractions. The remaining 559 cells presented very similar alignment rates, which were potential doublets. Among the 4,733 *R. breviuscula* cells, those whose alignment rates were lower than 25% were discarded, leaving 4,392 cells from *R. breviuscula* available for the next stage of the analysis.

### SNP calling and selection of markers in gametes

SNP calling in all gametes adopted the same methods as the reference genome SNP calling, e.g., via ‘bcftools mpileup’ and ‘bcftools call’ (v1.9), with the difference that the “--variants-only” flag was not applied. After acquiring SNPs for every gamete, the SNP positions, allele counts of the reference, and alternative bases were extracted through the ‘bcftools query’. Comparing SNPs in every gamete with markers defined on the reference resulted in reliable genotyping markers in this gamete.

Not all cells were suitable for CO calling due to insufficient markers or doublets generated during the 10X library construction; hence, filtering is necessary before CO alling. A total of 2,338 cells with fewer than 400 markers were first discarded to ensure accurate genotyping by sufficient markers. To remove doublets, the frequency of marker genotype switches across the remaining 2,054 cells was estimated. Cells with frequent switches, i.e., a switching rate (genotype switching times/number of markers) greater than 0.07, were taken as doublets (**Figure S4E**). Ultimately, 402 doublets were identified, with the remaining 1,652 cells proving suitable for subsequent CO calling.

### CO identification

The chromosome genotyping was performed by adapting the haplotype phasing method proposed by. The original approach was designed based on a scDNA-seq library, which is commonly used to examine more SNPs than scRNA-seq data; therefore, the smoothing function and parameters were adjusted to define the genotypes of genomic blocks accordingly. Specifically, the markers were first smoothed using neighbouring markers (two ahead and two behind) based on allele frequency and then on the presence of the genotypes. After smoothing, the genotype blocks containing at least five markers within 1 Mb were qualified to assign the genotypes. The genomic regions that saw the conversion of the genotypes at the flanks were taken as CO break positions (**Figure 4C,D**). Finally, the CO numbers in each cell were counted and manually assessed, and those with double COs were corrected.

### Recombination landscape and CO interference

To gain an overview of the CO rates across the chromosomes of *R. breviuscula*, the CO positions in all viable cells (1,641 cells remaining after manual correction) were summarised, and the recombination landscape for each chromosome was plotted (**Figure 5A**). Recombination rate (cM/Mb) was computed by 1-Mb sliding window and 100-kb step size.

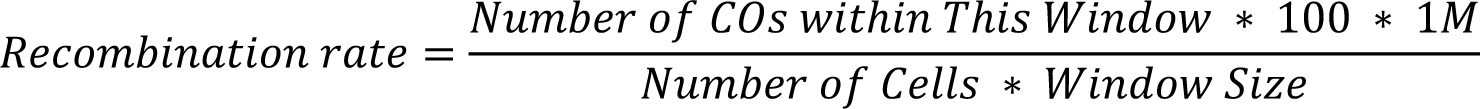

To plot the genetic linkage map (**Figure 5B**), 743 markers were extracted from the 820,601 reference markers by selecting the median marker within each 500-kb sliding window (step size was also 500 kb) from the first present marker until the last. CO interference was analysed with MADpattern (v.1.1)^76^, using 1,641 confident singleton pollen nuclei. Chromosome 1 was divided into 18 intervals and chromosomes 2–5 were divided into 15 intervals to compute the mean CoC of every pair of intervals.

### F1 offspring mapping and CO analysis

Sixty-three F1 offspring were reproduced from selfed *R. breviuscula*. Each F1 plant was sequenced with ∼3X Illumina WGS data. To genotype F1 offspring, WGS Illumina sequences of each plant were first mapped to rhyBreHap1 reference genome with bowtie2 (v2.4.4) paired-end mode, and then SNPs were called by ‘bcftools mpileup’ and ‘bcftools call’ (v1.9) (with --keep-alts, --variants-only, and --multiallelic-caller flags enabled). Next, SNPs of each F1 sample were input to TIGER^77^ for genotyping and generating potential CO positions. In addition, RTIGER^78^ was also used to identify the genotypes of chromosomal segments by utilizing the corrected markers resulted from TIGER. Only the COs that agreed by both tools were kept. The recombination landscape from F1 COs was plotted using the same strategy and sliding window as illustrated for pollen nuclei.

### ChIP

CENH3 ChIP-seq data were obtained from Hofstatter et al. ^32^. Further ChIP experiments were performed for H3K4me3 (rabbit polyclonal to Histone H3 tri-methyl K4; Abcam ab8580), H3K9me2 (mouse monoclonal to Histone H3 di-methyl K9, Abcam ab1220), H3K27me3 (mouse monoclonal to Histone H3 tri-methyl K27, Abcam ab6002), and the IgG control (recombinant rabbit IgG, monoclonal Abcam ab172730) using the same protocol described by Hofstatter et al. ^32^.

### ChIP-seq and analysis

ChIP DNA was quality-controlled using the next-generation sequencing assay on a FEMTO pulse (Agilent Technologies). An Illumina-compatible library was prepared with the Ovation Ultralow V2 DNA-Seq library preparation kit (Tecan Genomics) and sequenced as single-end 150-bp reads on a NextSeq2000 (Illumina) instrument. For each library, an average of 20 million reads were obtained.

Raw sequencing reads were trimmed using Cutadapt^79^ to remove low-quality nucleotides (with a quality score less than 30) and the adapters. Trimmed ChIPed 150-bp single-end reads were mapped to their respective reference genome using bowtie2 ^71^ with default parameters. All read duplicates were removed and only the single best matching read was kept on the final alignment BAM file. The BAM files were converted into BIGWIG coverage tracks using the bamCompare tool from deeptools^80^. The coverage was calculated as the number of reads per 50-bp bin and normalised as reads per kilobase per million mapped reads (RPKM). The magnified chromosome regions showing multiple tracks presented in **Figure 7B** were plotted with pyGenomeTracks^81^.

### *Tyba* array and CENH3 domain annotation

*Tyba* repeats were annotated using a BLAST search with a consensus *Tyba* sequence, allowing a minimum of 70% similarity. Further annotation of the *Tyba* arrays was performed by removing spurious low-quality *Tyba* monomer annotations shorter than 500 bp. Bedtools^82^ was used to merge all adjacent *Tyba* monomers situated at a maximum distance of 25 kb into individual annotations to eliminate the gaps that arise because of fragmented *Tyba* arrays, and those smaller than 2 kb were discarded.

CENH3 peaks were called with MACS3^83^ using the broad peak calling mode:

`macs3 callpeak -t ChIP.bam -c Control.bam --broad -g 380000000 --broad-cutoff 0.1`

The identified peaks were further merged using a stepwise progressive merging approach. CENH3 domains were generated by 1) merging CENH3 peaks with a spacing distance less than 25 kb using bedtools to eliminate the gaps that arise because of fragmented *Tyba* arrays or due to the insertion of TEs; and 2) removing CENH3 domains less than 1 kb in size.

### Transposable element annotation

Transposable element protein domains and complete LTR retrotransposons were annotated in the reference haplotype genome using the REXdb database (Viridiplantae_version_3.0)^84^ and the DANTE tool available from the RepeatExplorer2 Galaxy portal^85^.

### Enzymatic methyl-seq and analysis

To investigate the methylome space in *R. breviuscula*, the relatively non-destructive NEBNext Enzymatic Methyl-seq Kit was employed to prepare an Illumina-compatible library, followed by paired-end sequencing (2 × 150 bp) on a NextSeq2000 (Illumina) instrument. For each library, 10 Gb of reads was generated.

Enzymatic methyl-seq data were analysed using the Bismarck pipeline ^86^ following the standard pipeline described at https://rawgit.com/FelixKrueger/Bismark/master/Docs/Bismark_User_Guide.html. Individual methylation context files for CpG, CHG and CHH were converted into BIGWIG format and used as input tracks for the overall genome-wide DNA methylation visualisation with pyGenomeTracks and R plots.

### Quantitative correlation of COs and (epi)genetic features

The distribution and accumulation of all the different classes of (epi)genetic features were correlated with the distribution of the COs. Correlation matrix (**Figure 6B**) was calculated by Pearson correlation coefficient for each pair of all features under a 1-Mb smoothing window and 250-kb step size: specifically, mean CO rates, mean GC contents, CENH3 peak density, *Tyba* array density, SNP density, TE density, H3K4me3 RPKM, H3K9me2 RPKM, H3K27me3 RPKM, mean CpG, mean CHG, and mean CHH.

To inspect a possible centromere effect on CO positioning, the relative distance from the CO site was calculated to the closest left and right centromeric unit, i.e., the CENH3 domain or *Tyba* array, across the 378 COs in the F_1_ offspring and normalised all distances to 0–1 such that all neighbouring centromeric units were displayed in the same scale (**Figure 7G**). Crossover and marker positions over the transcript bodies, CENH3 domain or *Tyba* array were normalised by their distance to start sites and end sites and then counted by binning (**Figure 7E,F**).

To see the association of CO designations with a variety of (epi)genetic features at a local scale, we first counted the number of COs that overlap with CENH3, *Tyba* arrays, genes, TEs, LTRs, H3K4me3 peaks, H3K9me2 peaks, and H3K27me3 peaks by ‘bedtools intersect’ (v2.29.0). Next, we assigned 378 COs genome-wide at random. The number of COs on each chromosome was the same as that was detected by F1 individuals (e.g., 72 COs on chr1, 69 on chr2, 76 on chr3, 84 on chr4, and 77 on chr5), while the CO break gap length was picked up from the 378 real F1 CO gaps randomly. For each simulation round, the pseudo-COs were overlapped with (epi)genetic features again with ‘bedtools intersect’. Five thousand of these simulations were done, and the results were then plotted as the distribution of overlapped CO numbers for each feature (**Figure S10**). Finally, to evaluate the deviation of real overlapped COs with each feature to the expected overlapped CO number under the hypothesis of randomly distributed COs, Z-scores were calculated by the mean values and standard deviations of the simulated number of overlapped CO distribution (**Figure 7D**).

### Gene annotation

Structural gene annotation was done combining de novo gene calling and homology-based approaches with *Rhynchospora* RNAseq, IsoSeq, and protein datasets already available^32^.

Using evidence derived from expression data, RNAseq data were first mapped using STAR^87^ (version 2.7.8a) and subsequently assembled into transcripts by StringTie^88^ (version 2.1.5, parameters -m 150-t -f 0.3). Triticeae protein sequences from available public datasets (UniProt, https://www.uniprot.org, 05/10/2016) were aligned against the genome sequence using GenomeThreader^89^ (version 1.7.1; arguments - startcodon -finalstopcodon -species rice -gcmincoverage 70 -prseedlength 7 –prhdist 4). Isoseq datasets were aligned to the genome assembly using GMAP^90^ (version 2018-07-04). All assembled transcripts from RNAseq, IsoSeq, and aligned protein sequences were combined using Cuffcompare^91^ (version 2.2.1) and subsequently merged with StringTie (version 2.1.5, parameters --merge -m150) into a pool of candidate transcripts. TransDecoder (version 5.5.0; http://transdecoder.github.io) was used to identify potential open reading frames and to predict protein sequences within the candidate transcript set.

Ab initio annotation was initially done using Augustus^92^ (version 3.3.3). GeneMark^93^ (version 4.35) was additionally employed to further improve structural gene annotation. To avoid potential over-prediction, we generated guiding hints using the above described RNAseq, protein, and IsoSeq datasets as described by Nachtweide and Stanke^92^ A specific Augustus model for *Rhynchospora* was built by generating a set of gene models with full support from RNAseq and IsoSeq. Augustus was trained and optimized using the steps detailed by Nachtweide and Stanke^92^.

All structural gene annotations were joined using EVidenceModeller^94^ (version 1.1.1), and weights were adjusted according to the input source: ab initio (Augustus: 5, GeneMark: 2), homology-based (10). Additionally, two rounds of PASA^95^ (version 2.4.1) were run to identify untranslated regions and isoforms using the above described IsoSeq datasets.

We used DIAMOND^96^ (v2.0.5) to compare potential protein sequences with a trusted set of reference proteins (Uniprot Magnoliophyta, reviewed/Swissprot, downloaded on 3 Aug 2016; https://www.uniprot.org). This differentiated candidates into complete and valid genes, non-coding transcripts, pseudogenes, and transposable elements. In addition, we used PTREP (Release 19; https://trep-db.uzh.ch), a database of hypothetical proteins containing deduced amino acid sequences in which internal frameshifts have been removed in many cases. This step is particularly useful for the identification of divergent transposable elements with no significant similarity at the DNA level. Best hits were selected for each predicted protein from each of the three databases. Only hits with an e-value below 10e–10 were considered. Furthermore, functional annotation of all predicted protein sequences was done using the AHRD pipeline (https://github.com/groupschoof/AHRD).

Proteins were further classified into two confidence classes: high and low. Hits with subject coverage (for protein references) or query coverage (transposon database) above 80% were considered significant and protein sequences were classified as high-confidence using the following criteria: protein sequence was complete and had a subject and query coverage above the threshold in the UniMag database or no hit in UniMag but in UniPoa and not PTREP; a low-confidence protein sequence was incomplete and had a hit in the UniMag or UniPoa database but not in PTREP. Alternatively, it had no hit in UniMag, UniPoa, or PTREP, but the protein sequence was complete. In a second refinement step, low-confidence proteins with an AHRD-score of 3* were promoted to high-confidence.

BUSCO^97^ (version 5.1.2.) was used to evaluate the gene space completeness of the pseudomolecule assembly and structural gene annotation with the ‘viridiplantae_odb10’ database containing 425 single-copy genes.

### Ka/Ks ratio calculation

We identified homologs between *Brachypodium distachyon* (v3.0) (downloaded from ensemble plants plants.ensembl.org) and *Juncus effesus*^32^ using the ortholog module from JCVI python library^98^. Subsequently, pairwise alignments were generated with ParaAT^99^ (v2) and the Ka/Ks ratio was calculated using KaKs_Calculator^100^ (v3) using the YN method^101^. Plots were generated using karyoploteR^102^.

## Supplementary Information

**Table S1.** Summary of genome size, contigs and scaffolds of the phased genome assemblies.

|  | Haplotype 1 | Haplotype 2 |
| --- | --- | --- |
| **Genome assembly size (bp)** | 418,627,160 | 390,890,803 |
| **# Contigs** | 1637 | 548 |
| **Contig assembly size (bp)** | 421,256,472 | 391,742,506 |
| **Largest contig (bp)** | 35,313,519 | 43,961,622 |
| **Contig N50 (bp)** | 11,938,939 | 13,764,201 |
| **Contig N90 (bp)** | 42,248 | 2,739,863 |
| **# Scaffolds** | 1,501 | 457 |
| **Pseudo-chromosome size (bp)** | 368,174,147 | 370,478,156 |
| **Scaffold N50 (bp)** | 69,585,868 | 72,168,595 |
| **Scaffold N90 (bp)** | 45,843 | 66,381,717 |
| **Largest scaffold / chr 1 (bp)** | 91,632,052 | 89,220,796 |
| **Chromosome 2 (bp)** | 70,953,004 | 72,168,595 |
| **Chromosome 3 (bp)** | 69,585,868 | 69,956,709 |
| **Chromosome 4 (bp)** | 66,447,897 | 66,381,717 |
| **Chromosome 5 (bp)** | 69,555,326 | 72,750,339 |
| **Base accuracy (QV)** | 30.85 | 32.32 |
| **Completeness (%)** | 85 | 85 |
| **GC (%)** | 35.91 | 35.60 |

**Table S2.** Synteny and structural variations between two haplotypes of *R. breviuscula*.

|  |  |  |  |
| --- | --- | --- | --- |
| #Structural annotations |  |  |  |
| #Variation_type | Count | Length hap1 | Length hap2 |
| **Syntenic regions** | 229 | 329,130,991 | 329,924,075 |
| **Inversions** | 39 | 2,135,010 | 1,947,035 |
| **Translocations** | 346 | 3,620,755 | 3,569,626 |
| **Duplications (reference)** | 137 | 1,472,557 | – |
| **Duplications (query)** | 249 | – | 1168366 |
| **Not aligned (reference)** | 650 | 32,783,105 | – |
| **Not aligned (query)** | 808 | – | 33606738 |
| #Sequence annotations |  |  |  |
| #Variation_type | Count | Length hap1 | Length hap2 |
| **SNPs** | 615,883 | 615883 | 615,883 |
| **Insertions** | 59,142 | – | 2,687,428 |
| **Deletions** | 59,276 | 3,101,459 | – |
| **Copy gains** | 87 | – | 126,950 |
| **Copy losses** | 60 | 394,961 | – |
| **Highly diverged** | 5,660 | 172,894,800 | 174,131,686 |
| **Tandem repeats** | 3 | 482 | 825 |

**Figure S1.**
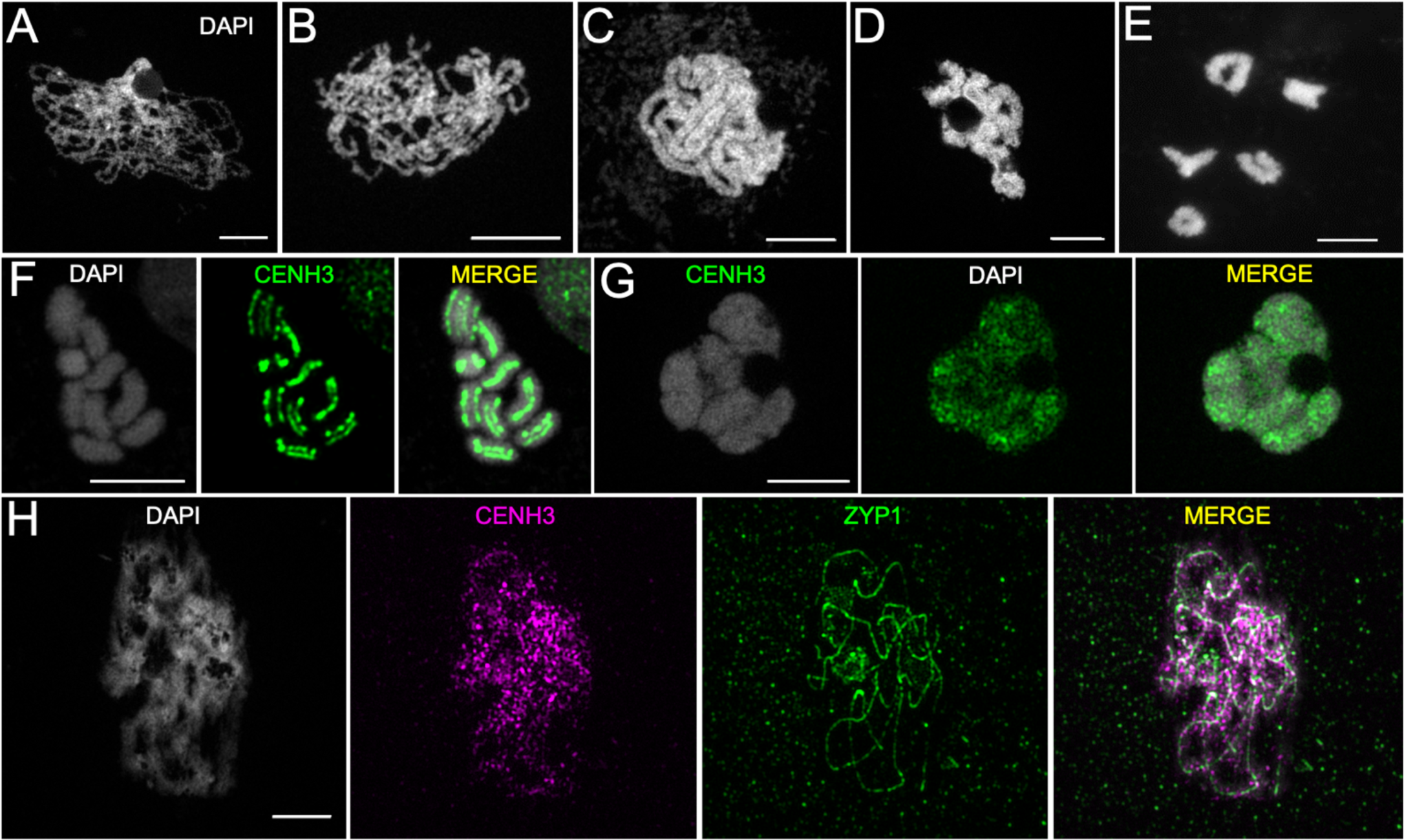
Chromosome spreads and immunolocalisation in male *R. breviuscul*a meiocytes.

**Figure S2.**
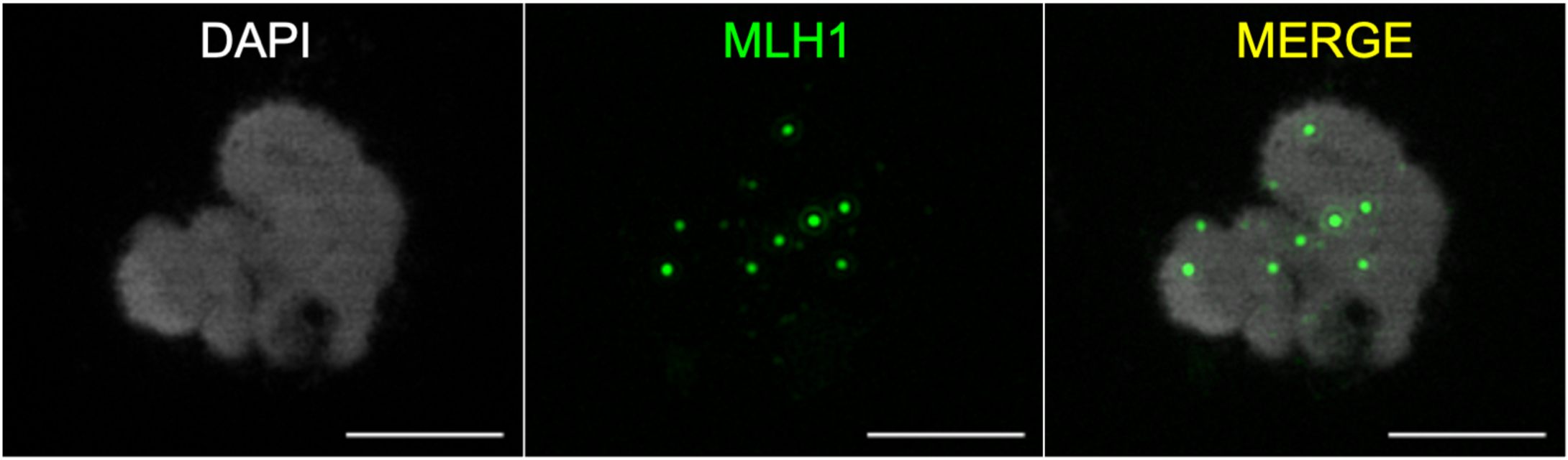
Maximum number of MLH1 (green) foci observed in *R. breviuscula* at diplotene.

**Figure S3.**
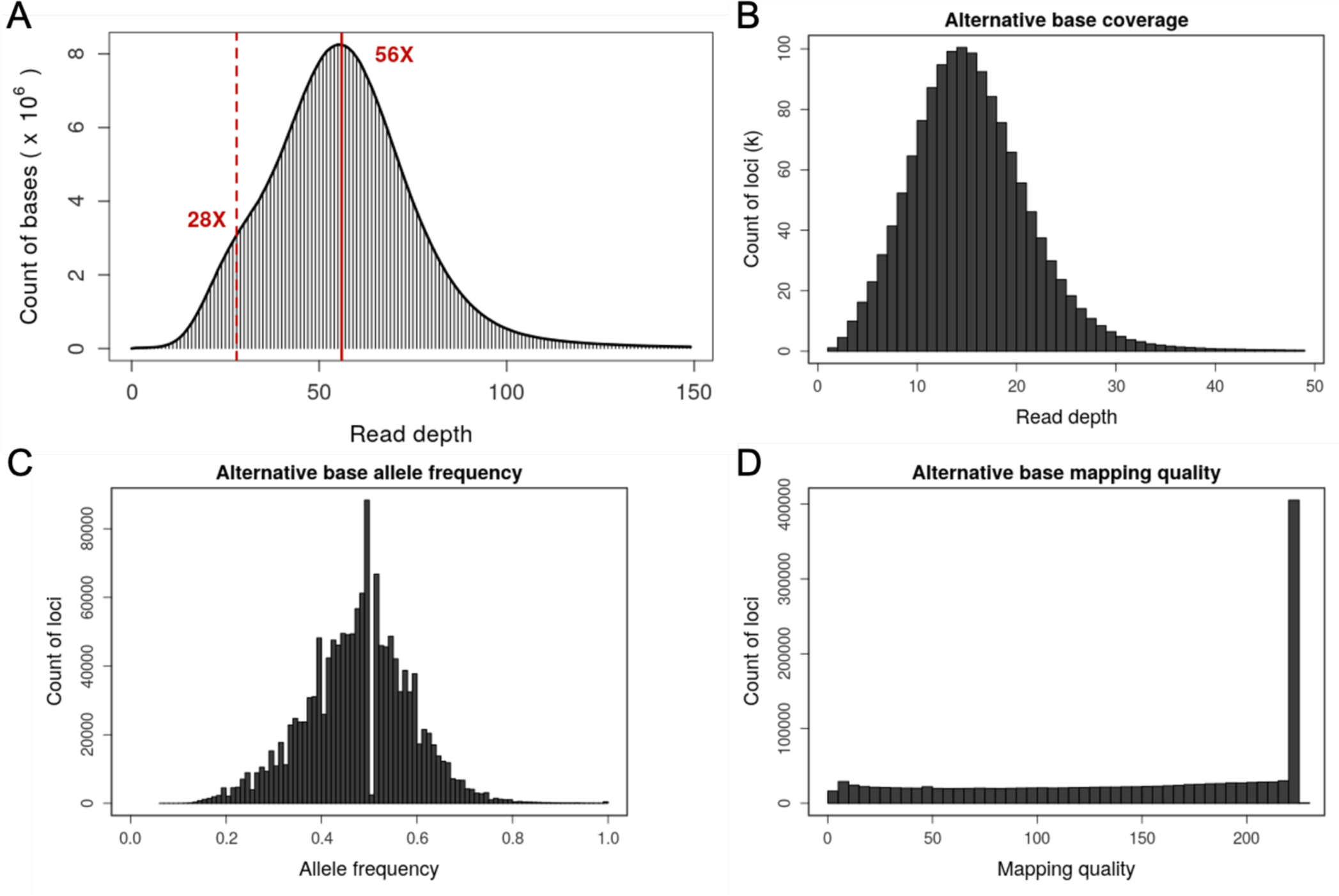
Selection of genotyping markers on the reference genome.

**Figure S4.**
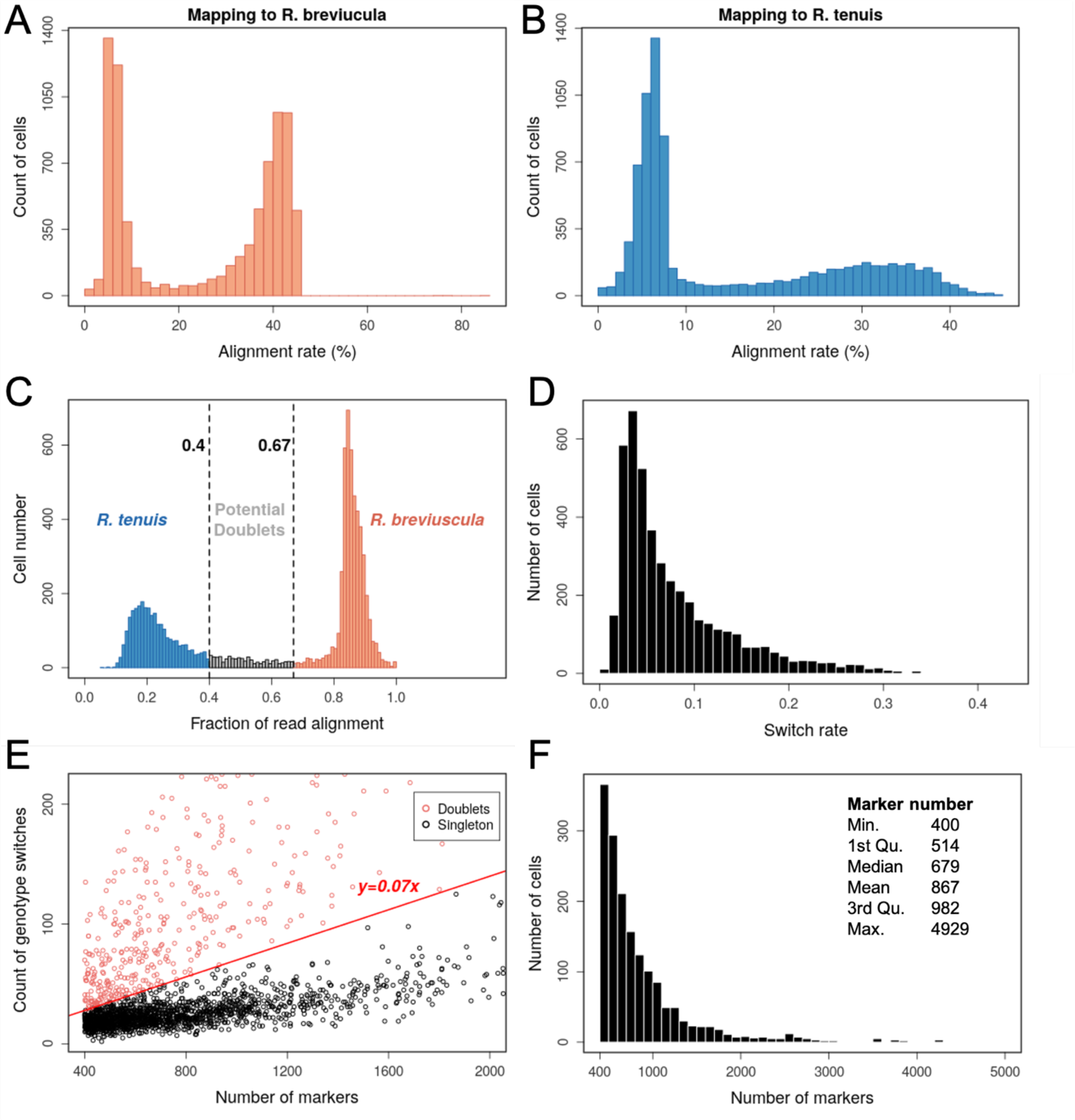
Pre-processing of scRNA-seq by separating *R. breviuscula* from *R. tenuis* cells and removing doublets.

**Figure S5.**
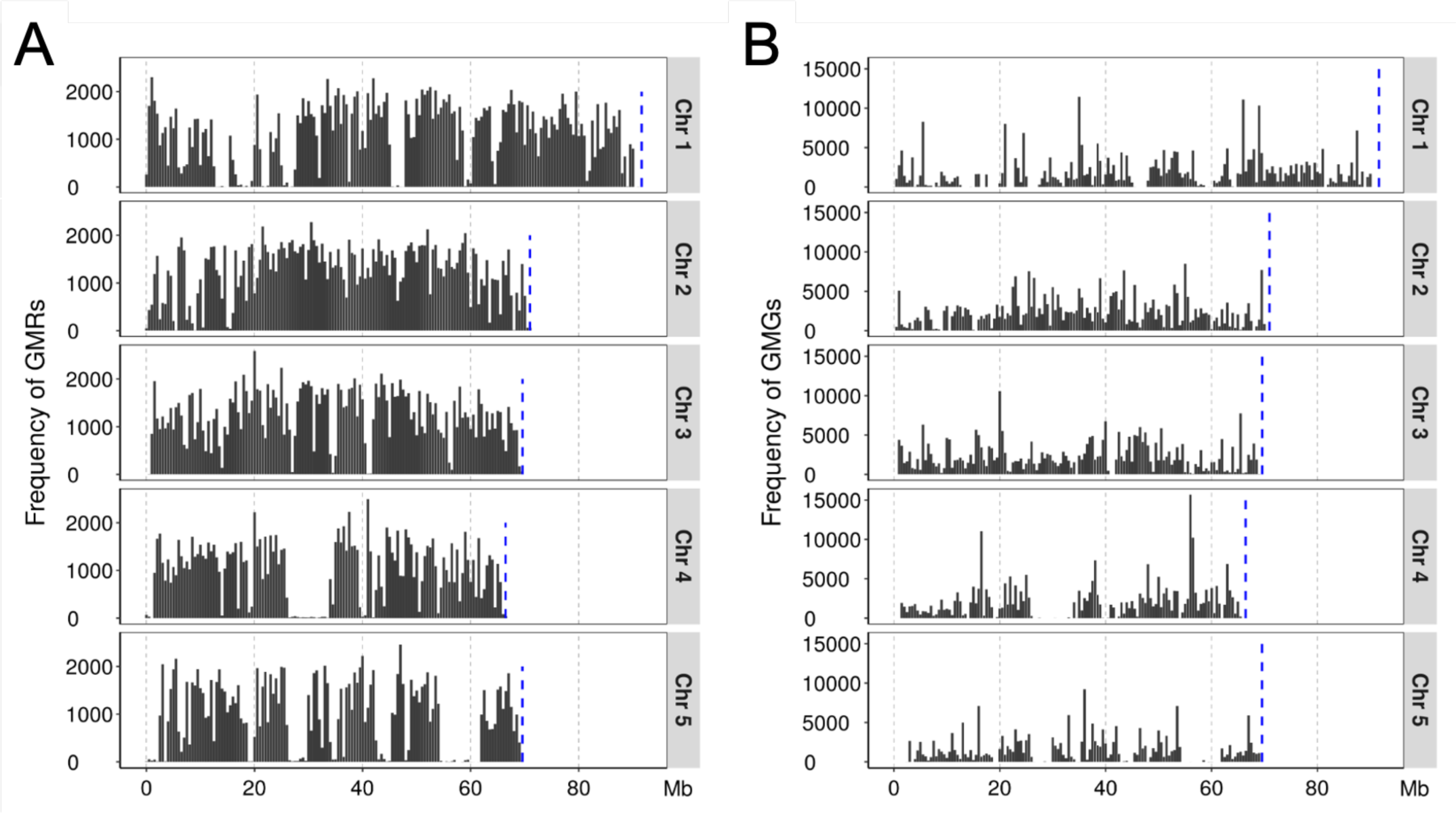
Marker distribution on the reference and across all viable pollen nuclei.

**Figure S6.**
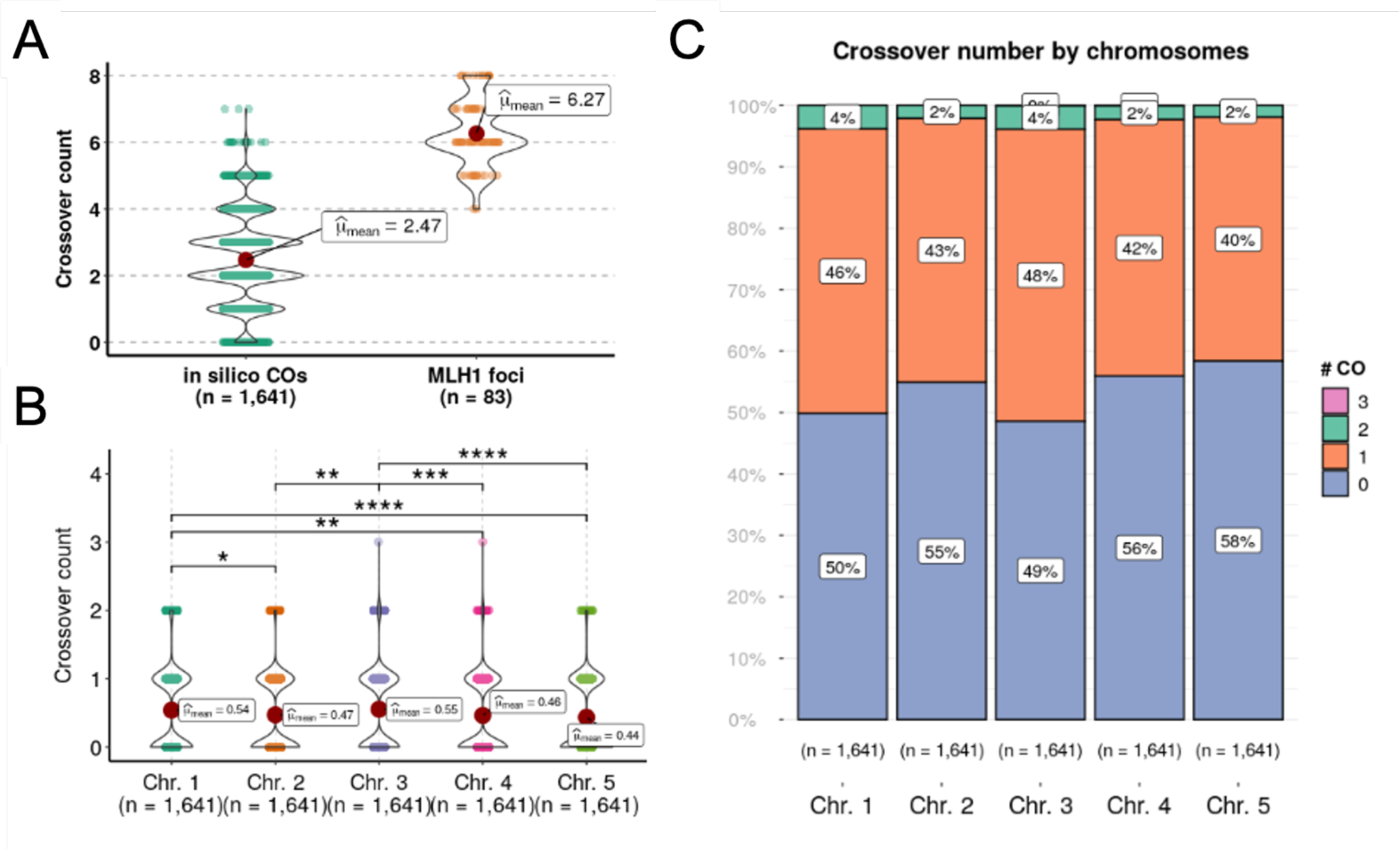
Number of COs in all viable pollens.

**Figure S7.**
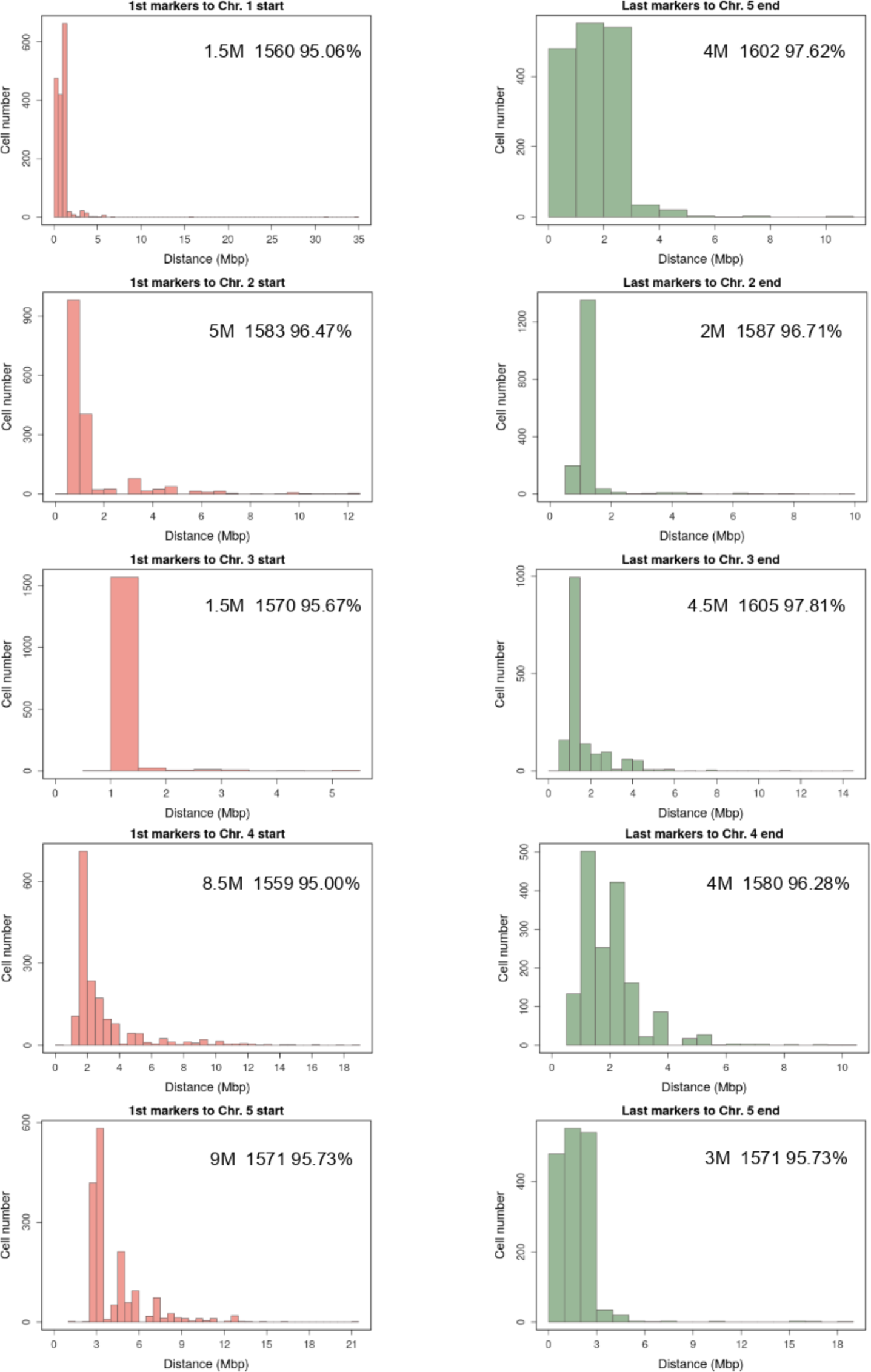
Distance distribution of the first markers to the chromosome start and the last markers to chromosome ends across all viable pollen nuclei.

**Figure S8.**
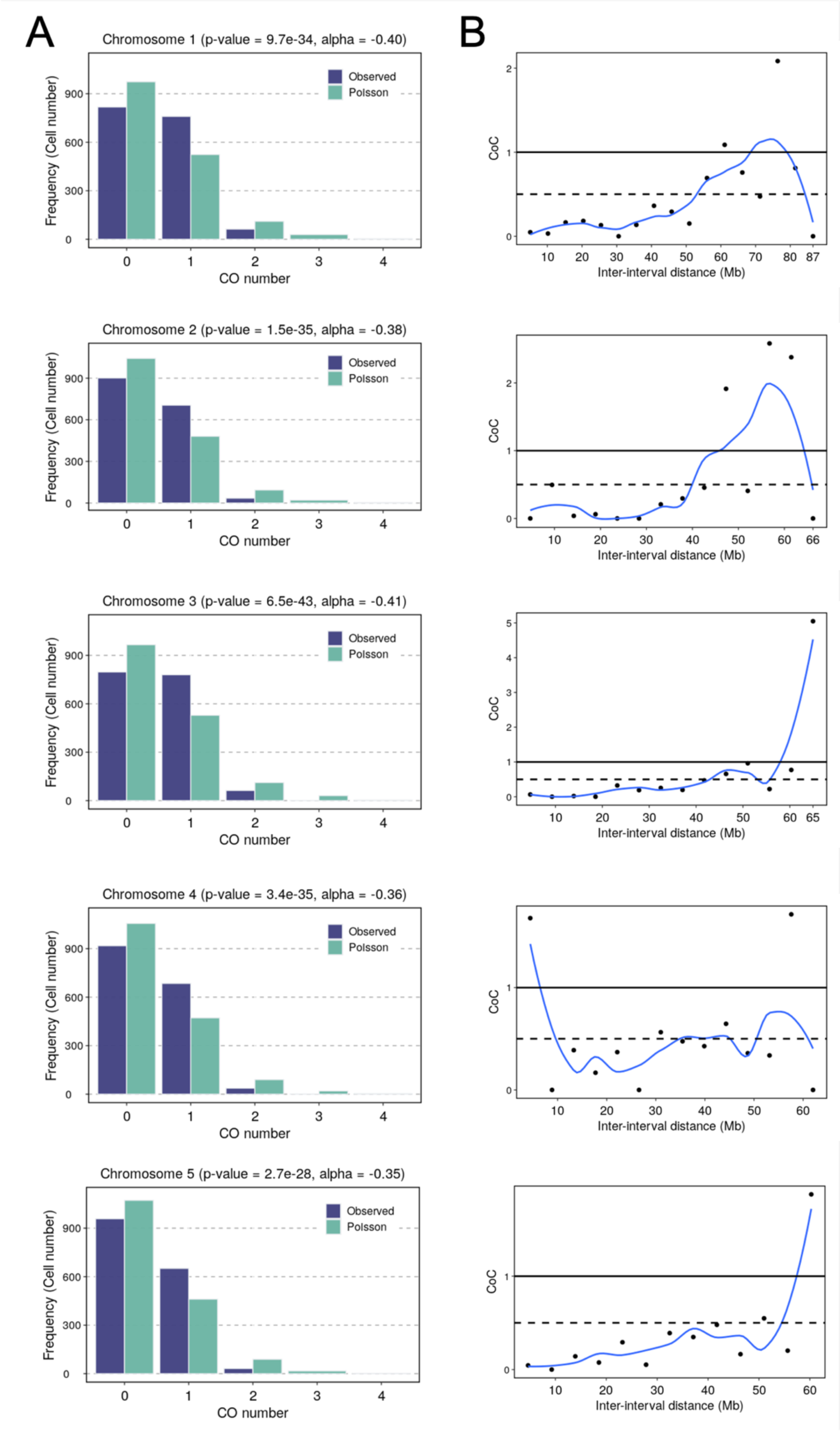
CO interference on CO number.

**Extended Data Fig 9.**
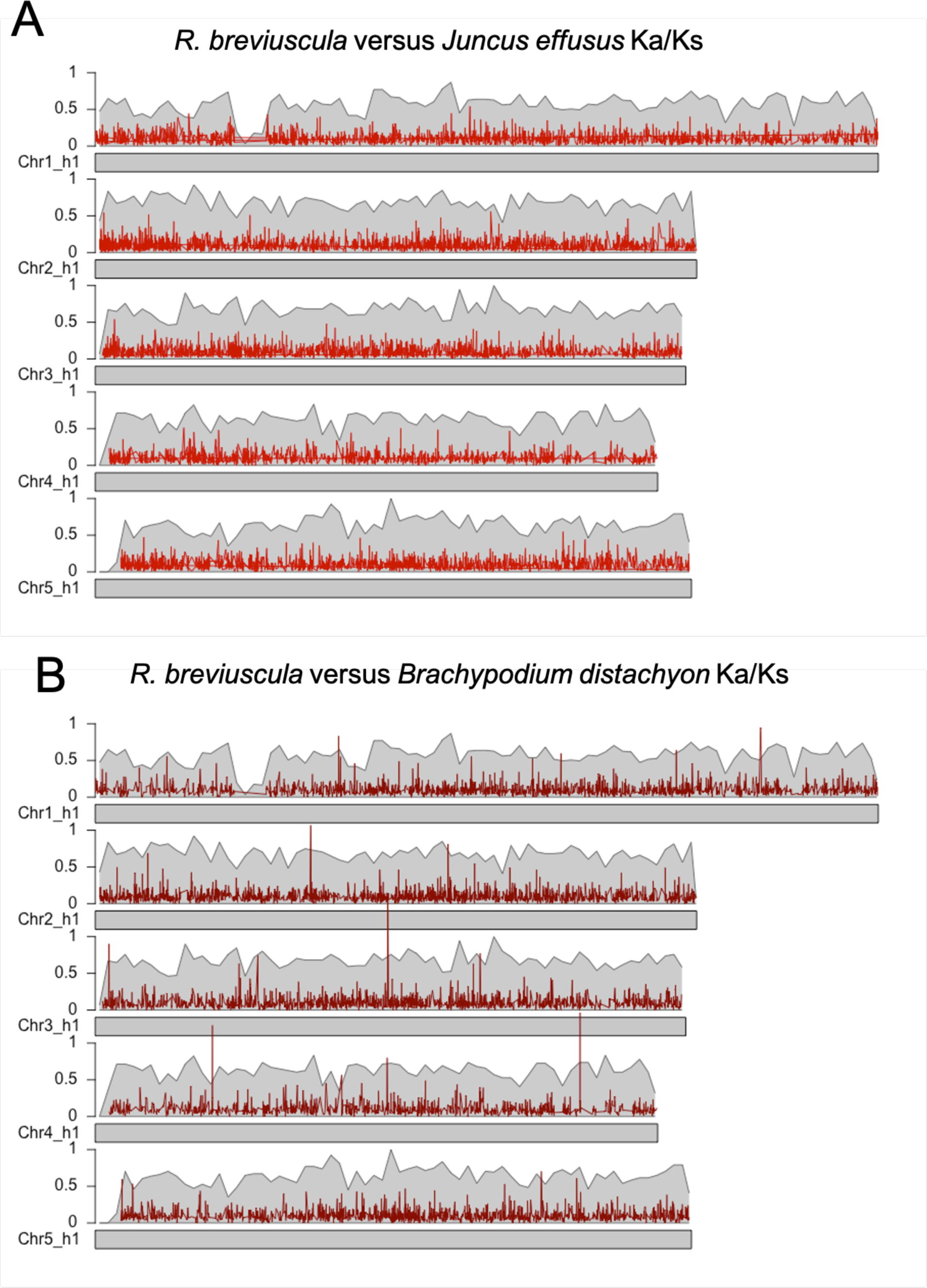
Ka/Ks ratio estimation across the chromosomes of *R. breviuscula*.

**Figure S10.**
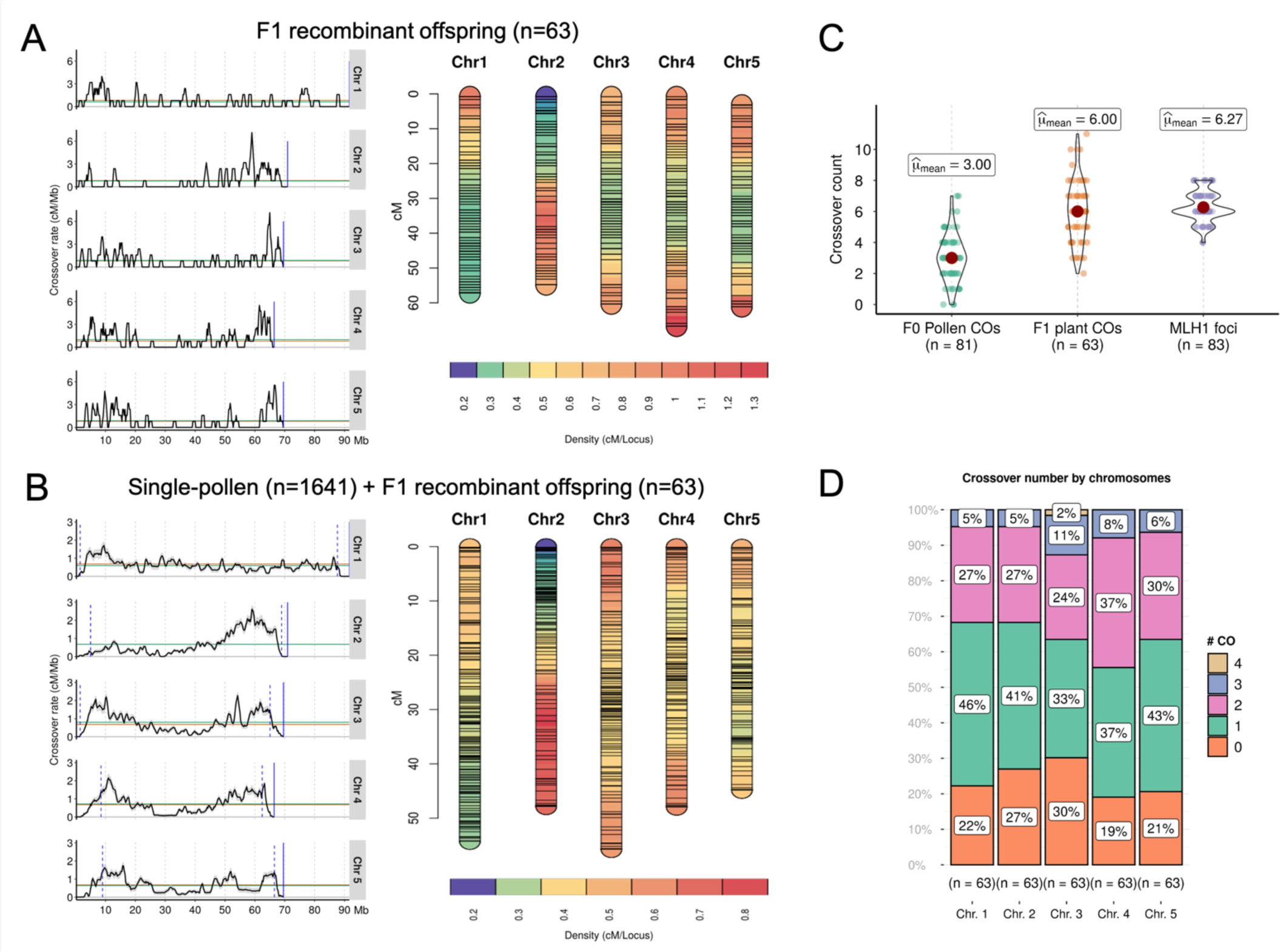
Recombination dynamics in the F1 recombinant offspring and combined data (F1 + le-pollen sequencing) of *R. breviuscula*.

**Figure S11.**
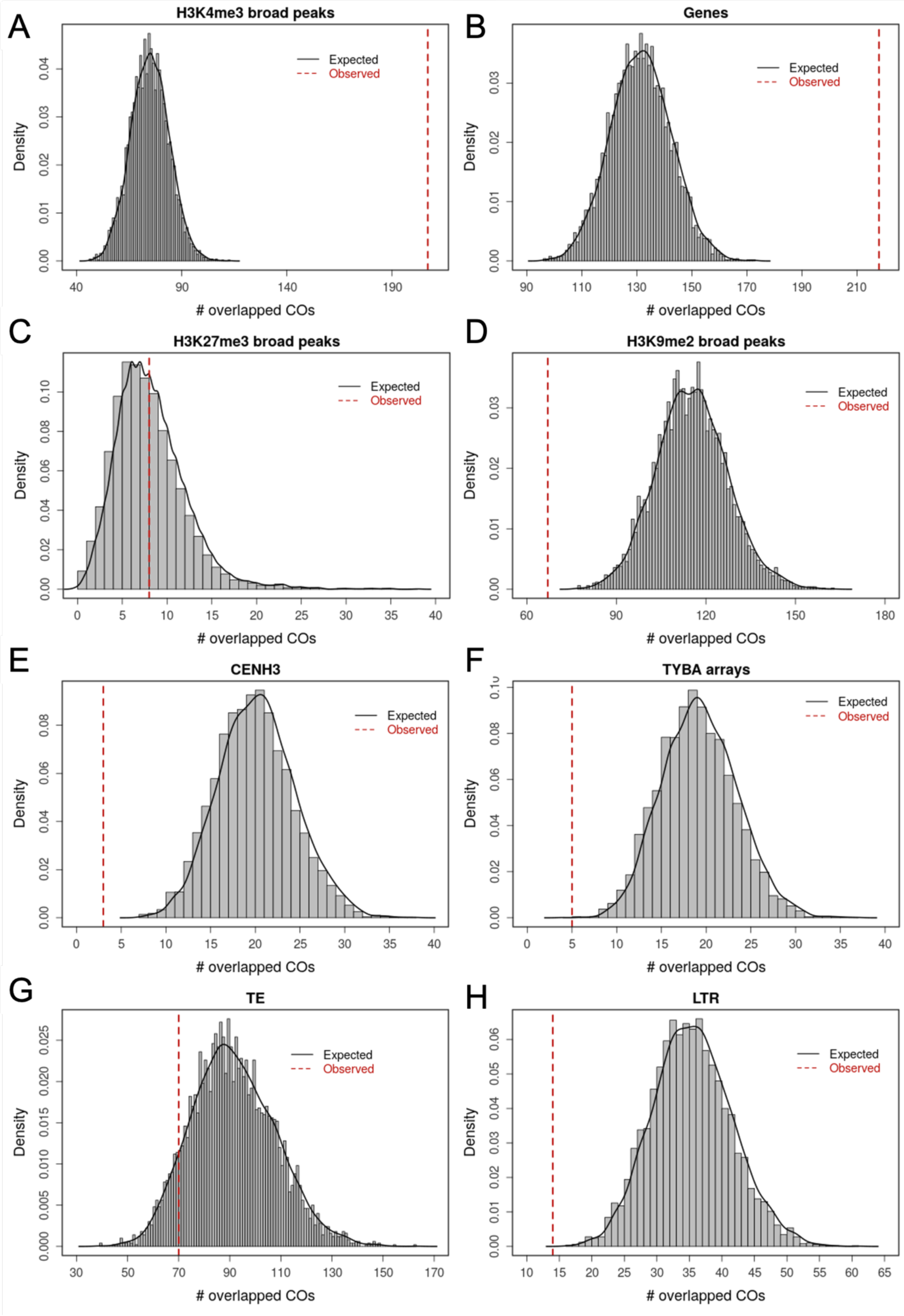
Comparison of numbers of COs overlapped with (epi)genetic features to random simulations.

**Figure S12.**
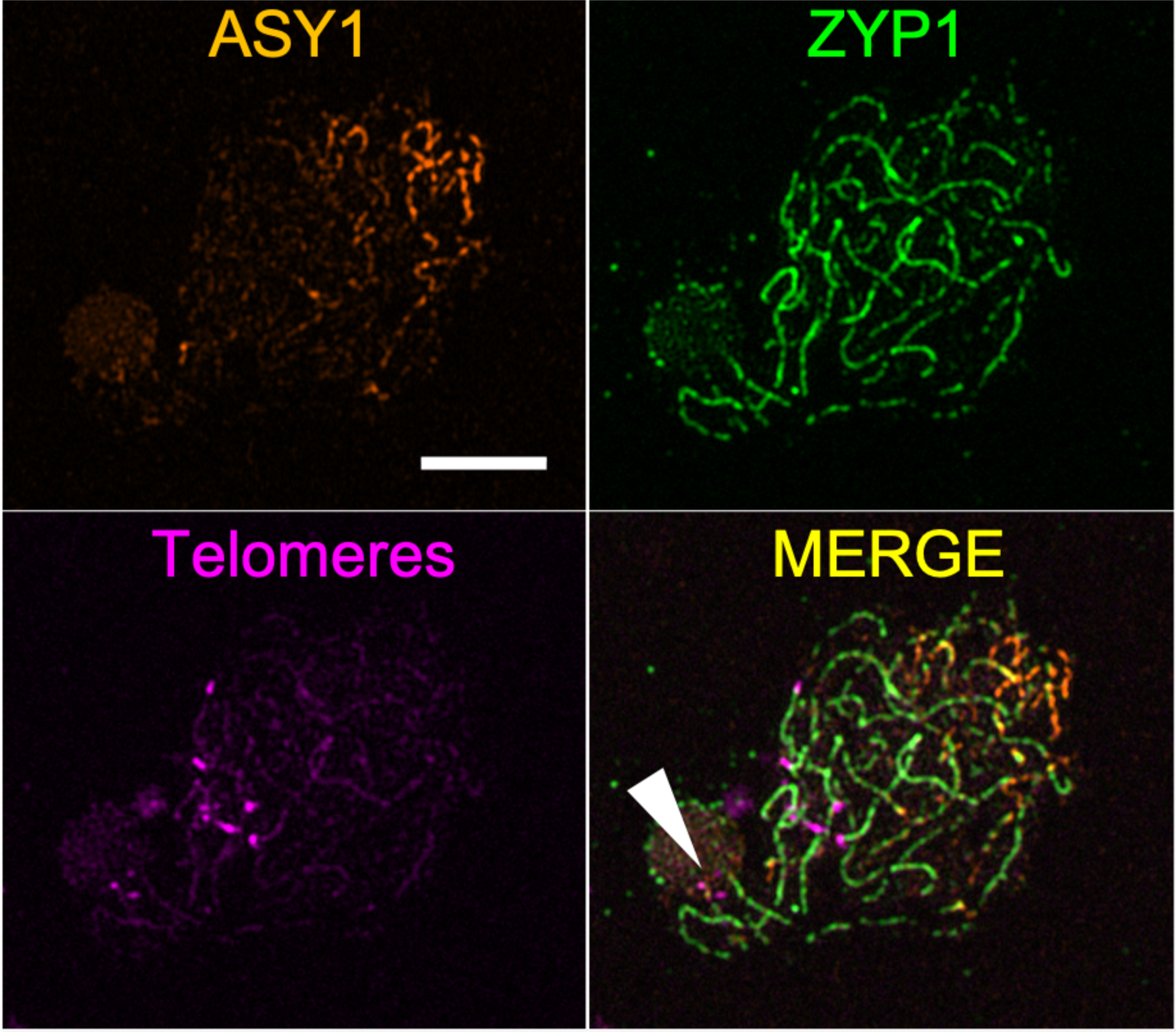
Immunolocalisation of ASY1, ZYP1 and telomere-FISH.

**Figure S13.**
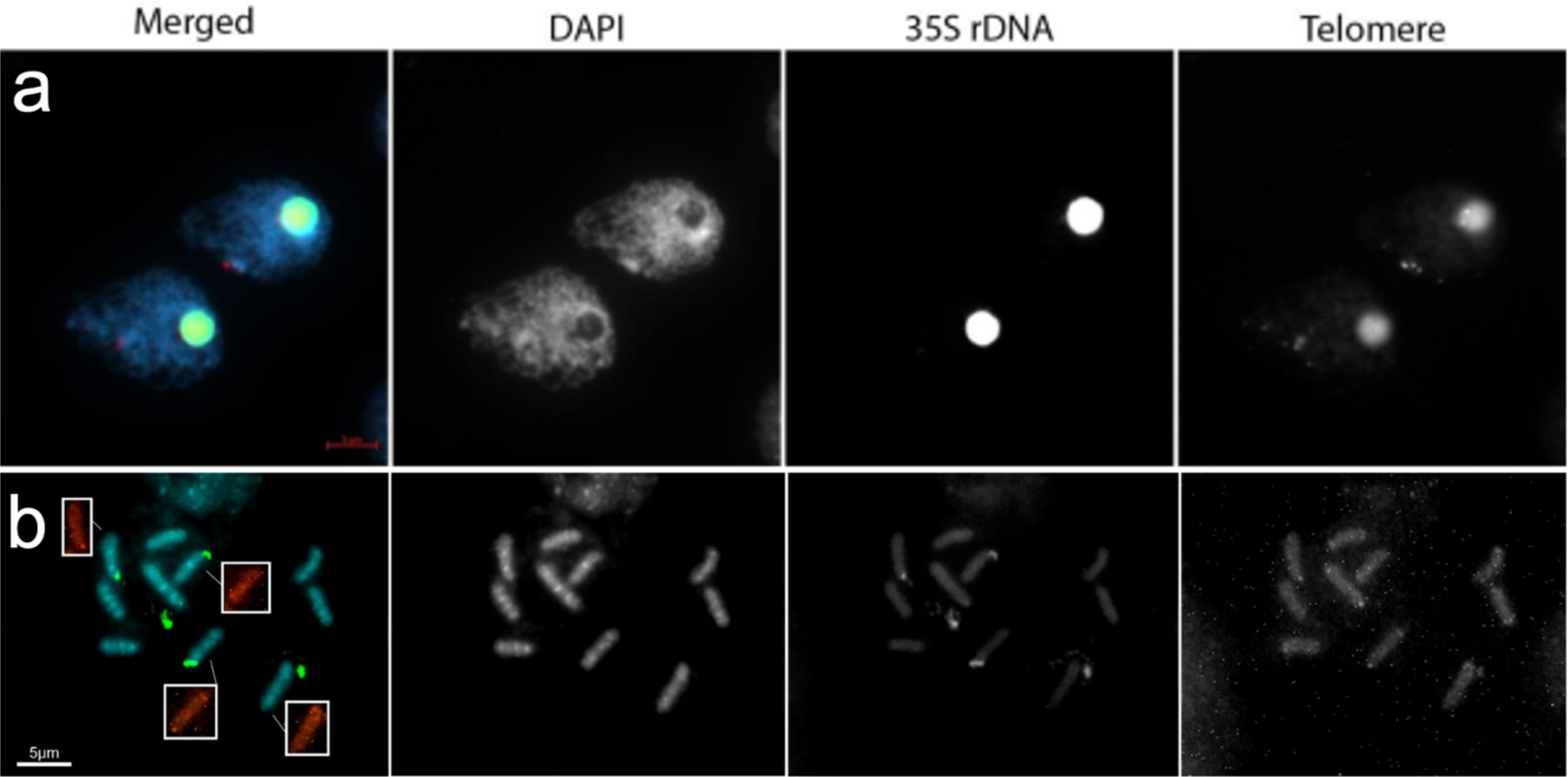
FISH with 35S rDNA and a telomeric probe in *R. breviuscula*.

